# Targeting a critical step in fungal hexosamine biosynthesis

**DOI:** 10.1101/2020.01.07.896944

**Authors:** Deborah E.A. Lockhart, Mathew Stanley, Olawale G. Raimi, David A. Robinson, Dominika Boldovjakova, Daniel R. Squair, Andrew T. Ferenbach, Wenxia Fang, Daan M.F. van Aalten

**Author notes:** These authors contributed equally to this work (Deborah E.A. Lockhart, Mathew Stanley).

## Abstract

*Aspergillus fumigatus* is a human opportunistic fungal pathogen with a cell wall that protects it from the extracellular environment. Chitin, an essential cell wall component, is synthesised from UDP-GlcNAc that is produced by the hexosamine biosynthetic pathway. Here, we provide genetic and chemical evidence that glucosamine 6-phosphate *N*-acetyltransferase (Gna1), a key enzyme in this pathway, is an exploitable antifungal drug target. Deletion of *GNA1* results in loss of viability and disruption of the cell wall, phenotypes that can be rescued by the product of the enzyme. In a murine model of aspergillosis, the *Δgna1* mutant strain attenuates virulence. Using a fragment-based approach, we discovered a small heterocyclic scaffold that binds proximal to the active site and can be optimised to a selective sub-micromolar binder. Taken together, we have provided genetic, structural and chemical evidence for Gna1 as an antifungal target in *Aspergillus fumigatus*.

## Introduction

*Aspergillus* spp., *Candida* spp., and *Cryptococcus* spp. are opportunistic human pathogens that together account for most global invasive fungal infections. Considered a significant and growing challenge to human health^1,2^, infections attributable to *Aspergillus fumigatus* are a significant cause of morbidity and mortality in an ever-expanding group of patients. For patients undergoing haematopoietic stem cell transplantation (HSCT) and in particular allogeneic grafts, invasive aspergillosis (IA) is an important cause of pulmonary related mortality^3^. Chronic and allergic forms are rarely life threatening, however, they are estimated to have global burdens of approximately 1.2 million and 4.8 million people respectively^4,5^. Moreover, they may affect patients with an intact immune system but have other medical co-morbidities placing them at a higher risk of fungal sensitisation and subsequent allergic disease. A recent phenomenon in both immunosuppressed and immunocompetent individuals is post influenza aspergillosis in the Intensive Care Unit (ICU)^6^.

As both fungi and mammals are eukaryotes, antifungal drug development faces the fundamental challenge of selective toxicity. Structures that are unique to fungi or that bear exploitable dissimilarities, such as the cell membrane and the cell wall, are exploited by the polyenes^7^, azoles^8^ and echinocandins^9^ respectively. Together with flucytosine^10^, they represent the only antifungal classes clinically licensed for the prevention and treatment of aspergillosis^3^ and offer limited degrees of efficaciousness through a restricted spectrum of cellular targets, poor clinical and physicochemical properties and rising rates of drug resistance^11^ particularly exacerbated through extensive agricultural over-use of antifungal azoles^2,12^. A lack of innovation, investment and infrastructure in pharmaceutical antifungal research has been cited for the dearth of preclinical candidates despite a global market worth in excess of US $6 billion^13^. Traditionally antifungal agents were discovered by screening large libraries of natural products or synthetic small molecules for their fungistatic and, preferably, fungicidal properties^14^. The polyenes and echinocandins were both derived from natural product screening. Unfortunately many subsequent screens ‘rediscovered’ the same basic chemical scaffolds^14^. The antifungal pipeline has displayed modest progression^15^, with only a single new antifungal class currently undergoing late-stage clinical trials (Phase IIb, NCT03583164, ClinicalTrials.gov). While the introduction of additional azoles such as isavuconazole^16,17^ provides greater options to reduce side effects and simplify dosing regimens, the development of resistance is not adequately addressed by new members of this widely used class^18^.

A major characteristic feature of fungi is the presence of a highly ordered carbohydrate cell wall that provides protection against chemical and mechanical stresses. The cell wall is a dynamic, interlaced and only partially defined polysaccharide structure that is essential for survival^19^ and is considered an attractive source of novel antifungal targets due to its absence in humans. The *A. fumigatus* cell wall predominantly consists of a central core of fibrils composed of branched β-1,3 glucan cross-linked to chitin. Polysaccharides such as β-1,3-1,4-glucan and galactomannan are covalently bound to this complex while those contained in the surrounding cement include α-1-3 glucan. Cell wall biosynthesis and remodelling is a highly organised and complex process although many of the key steps remain to be fully elucidated^19^. Polysaccharide synthesis of β-1,3 glucan and chitin occur at the fungal cell membrane using intracellular sugar nucleotide donors as substrates. Chitin is an integral structural component of the cell wall in *A*. *fumigatus* contributing to rigidity and consists of a linear polymer of β(1-4)-linked *N*-acetyl-D-glucosamine (GlcNAc). Chitin synthesis remains an attractive drug target, however, targeting the entire chitin synthase family is not a viable option due to predicted differences in structure and the membrane bound location of the eight proteins^20^.

An alternative target to chitin synthases is biosynthesis of the sugar nucleotide substrate UDP-GlcNAc that is also utilised for the synthesis of GPI anchors and *N*- and *O*-linked glycans^21^. Four highly conserved enzymes in UDP-GlcNAc biosynthesis form the hexosamine biosynthetic pathway^22^ which is believed to represent the only endogenous source of UDP-GlcNAc within fungal cells^23^. Approximately 2-5% of glucose that enters the cell is directed into this pathway where it is phosphorylated to glucose-6-phosphate (Glc-6P) and subsequently converted to fructose-6-phosphate (Fruc-6P). Four enzymes catalyse the remaining steps of this pathway. Glucosamine-6-phosphate synthase (Gfa1) converts Fruc-6P to glucosamine-6-phosphate (GlcN-6P). This undergoes acetylation by glucosamine-6-phosphate *N*-acetyltransferase (Gna1) to *N*-acetyl-glucosamine-6-phosphate (GlcNAc-6P). The next step involves isomerisation by phospho-acetyl-glucosamine mutase (Agm1) to yield *N*-acetyl-glucosamine-1-phosphate (GlcNAc-1P) before uridylation by *N*-acetyl-glucosamine-1P-uridyltransferase (Uap1) to produce UDP-GlcNAc. This pathway is highly regulated in filamentous fungi, yeast and higher eukaryotes^23^. Three of the enzymes (Gfa1, Agm1 and Uap1) have been genetically validated in *A. fumigatus* as essential for growth under *in vitro* laboratory conditions^24–26^.

Gna1 (EC 2.3.1.4) is a member of the Gcn5-related *N*-acetyltransferase (GNAT) superfamily and is present in yeast such as *Candida* spp. (e.g. *Candida albicans*, *Candida glabrata* and *Candida auris*) and *Saccharomyces cerevisiae*. Genetic studies in *S. cerevisiae* have demonstrated that deletion of *GNA1* is lethal^27^. In *C*. *albicans GNA1* is essential for *in vitro* viability and attenuated virulence in a murine model of disseminated candidiasis^28,29^. Inactivation of the murine *GNA1* homologue is lethal as it is implicated in mitotic membrane fusion events and this further reinforces a critical role for this enzyme in higher eukaryotes^30^. Though evolutionarily and structurally distinct from higher eukaryotic organisms, the *GNA1* orthologues recently identified in parasitic protozoan apicomplexans have been suggested as potential therapeutic targets^31^.

To date, Gna1 in *A. fumigatus* is unexplored as an antifungal drug target using a combination of genetic and chemical methods. Inhibitors that chemically phenocopy a *GNA1* deletion have not been reported in any organism and the high degree of sequence homology between eukaryotic *GNA1* family members likely preclude fungal selectivity. Here, we report a multidisciplinary approach providing genetic, chemical and structural evidence supporting *A. fumigatus* Gna1 as an antifungal target and describe a fragment that exploits a previously unidentified pocket selective for the fungal enzyme.

## Results

### Disruption of gna1 leads to a terminal phenotype that is rescued by exogenous GlcNAc

To investigate the requirement for Gna1 in *A. fumigatus* UDP-GlcNAc biosynthesis, we deleted *GNA1* by homologous recombination. A *Δgna1* strain was constructed by replacing the coding sequence with a *pyrG* selection marker (Figure S1). Transformants were evaluated by phenotypic and PCR screening before verification by Southern Blot (Figure S1). Using a similar approach, *GNA1* (with a *N*-terminal HIS tag) was re-introduced into the *Δgna1* mutant (Figure S1). Although suitable for initial *in vitro* studies, it was imperative that the reconstituted strain restored *pyrG* prototrophy as *pyrG*-*A. fumigatus* strains are highly attenuated in murine models of aspergillosis^32^. To address this, the *pyrG*-strain was targeted with random integration of *A. fumigatus pyrG* into the chromosome to generate a *pyrG*+ reconstituted strain (Figure S1).

*In vitro* growth of *A. fumigatus Δgna1* was completely inhibited in glucose (Glc) supplemented media and restored to comparable levels upon replacement of glucose with GlcNAc, the product of the Gna1 enzyme (Fig. 1a). This demonstrates that *GNA1* is essential for viability of *A. fumigatus* in standard *in vitro* laboratory media rich in glucose and concurs with previous work in yeast^27,29^. We considered the physiological relevance of these Glc and GlcNAc concentrations since failure to correlate with physiological nutritional availability may incorrectly assign a gene as essential for growth. The glucose concentration used is 20-fold greater than the mean physiological human blood glucose level (110 mM vs. ∼5.5 mM) and in lung alveoli glucose concentrations are further reduced (∼ 0.1 mM)^33^. Specific human physiological levels of GlcNAc are unspecified in the literature but regarded “low” and subject to fluctuation. We initially investigated rescue of *Δgna1* on solid media supplemented with Glc (0.1 mM) and a simulated range of physiological GlcNAc concentrations, above and below previously reported levels^29^ (Fig. 1a). The *Δgna1* growth phenotype in the presence of “low” GlcNAc levels (1.5 μM) was characterised by a complete loss of hyphae and melanin pigment. “High” GlcNAc levels (150 μM, 100-fold increase) partially restored hyphal development but colonies were still strikingly devoid of pigmentation and overall radial growth was less than the parental and reconstituted strain. We describe these characteristics as a terminal phenotype.

**Figure 1.**
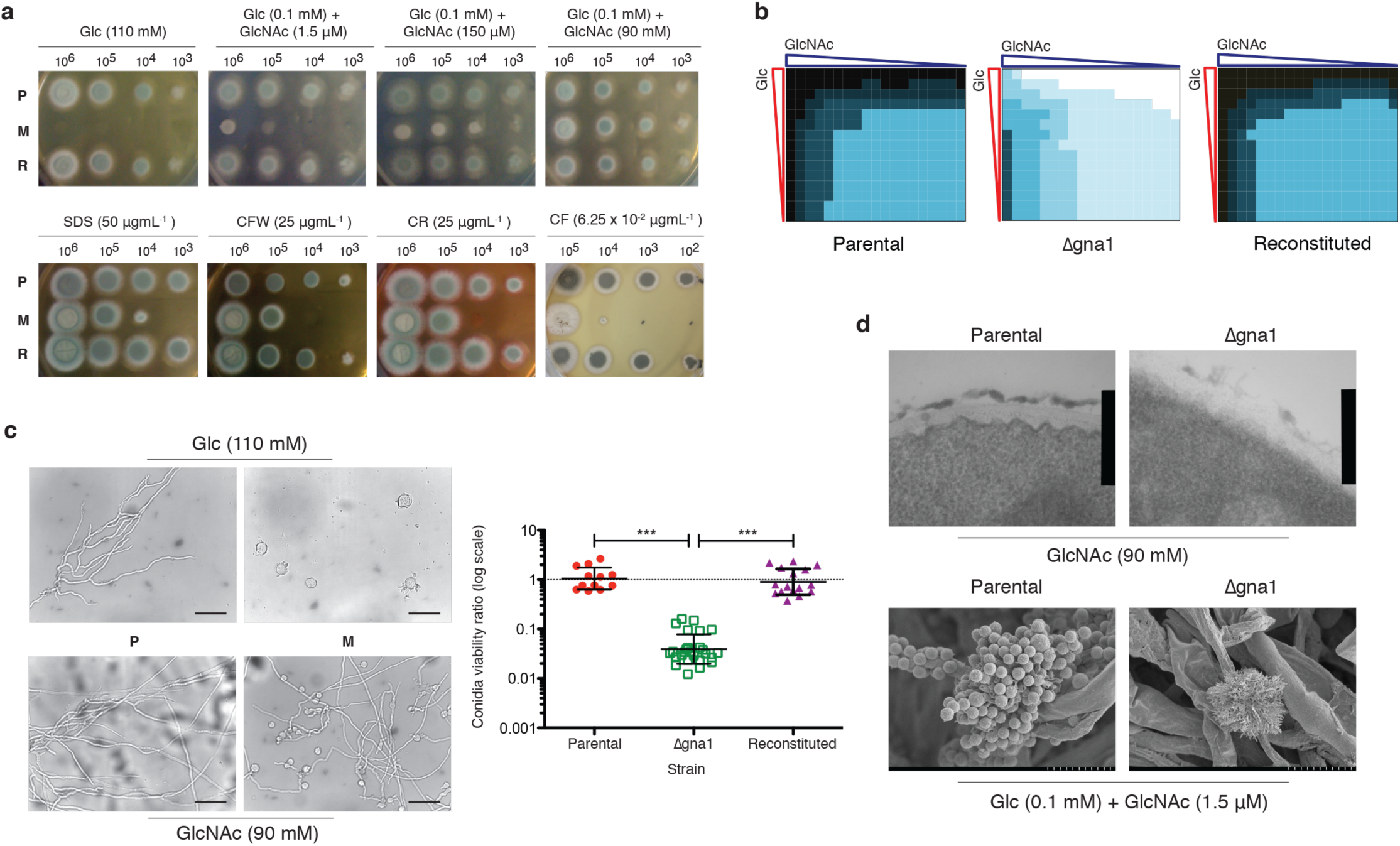
Disruption of *gna1* leads to a terminal cell wall phenotype partially rescued by GlcNAc. **a** Upper panel, colony morphology after 24 h incubation following inoculation of viable *A. fumigatus* conidia on CM supplemented with Glc and GlcNAc (as indicated). The morphology of *Δgna1* in the central two images is indicative of the terminal phenotype. Lower panel, serial dilutions of viable conidia were inoculated onto solid CM (0.1 mM Glc + *≥* 50 mM GlcNAc) supplemented with Sodium Dodecyl Sulphate (SDS), Calcofluor White (CFW), Congo Red (CR), or Caspofungin (CF) and incubated for 48 h. Images are representative of results obtained from three independent experiments (*n* = 3). **b** Heatmap of growth after 48 h incubation (inoculum 1 × 10^3^ viable conidia) in a checkerboard assay containing combinations of Glc and GlcNAc from 0 to 50 mM (2-fold serial dilutions). The parental, *Δgna1* mutant and reconstituted strain were scored numerically from one (no growth / inoculum spot) to seven (confluent growth with dark pigmented conidia). Each square denotes an individual condition and represents the mean growth from three independent experiments (*n* = 3). Black (maximum growth) to white (no growth). **c** Exogenous Glc arrests conidial germination and GlcNAc rescue fails to fully restore viability of *Δgna1*. Left panel, conidia were inoculated into liquid CM containing Glc (upper) GlcNAc (lower), incubated for up to 24 h and examined using a Leica light microscope at ×40 magnification. Scale bar 0.05 mm. Right panel, ratio of viable conidia versus total number of conidia. Each data point represents an individual replicate from a minimum of three independent experiments; horizontal lines represent the mean ± SD (*n* = 3). A ratio of one denotes no difference. *** *P* < 0.0001. **d** Viable conidia (1 × 10^5^) were inoculated onto solid CM supplemented with Glc and GlcNAc and incubated for 16 h (hyphae, upper panel) or 96 h (conidia, lower panel). Samples were fixed, processed and examined using electron microscopy (upper panel, transmission electron microscopy [TEM]; lower panel, scanning electron microscopy [SEM]). Scale bar: upper panel, 361 nm; lower panel, 10 μm. P, parental; M, *Δgna1* mutant; R, reconstituted *A. fumigatus* strains. All incubations were at 37 °C.

To dissect the interplay between exogenous Glc and GlcNAc, we developed a checkerboard assay to assess 285 different nutritional combinations of these to explore possible concentrations encountered by *A. fumigatus* in the human host and environment (Fig. 1b). Heatmap analysis demonstrated that loss of *GNA1* is associated with a terminal phenotype defective in conidial pigmentation and hyphal growth across a range of estimated physiological concentrations. Taken together, these experiments show that disruption of *gna1* leads to a terminal phenotype that is rescued by exogenous GlcNAc.

### The Δgna1 strain displays a cell wall phenotype

Next, we investigated whether rescue of *GNA1* with exogenous GlcNAc induces a cell wall stress response. *A. fumigatus* harbours cell wall integrity pathways that are activated under conditions of stress by signalling pathways such as the mitogen activated protein kinase (MAPK) cascades^34^. Due to the terminal phenotype not yielding sufficient growth for comparative analyses, this experiment was performed using a GlcNAc concentration of 90 mM to rescue growth. At concentrations of ≤ 10^4^ viable conidia, the *Δgna1* mutant displayed increased sensitivity to agents compromising the cell wall and membrane, suggesting the presence of cell wall defects and/or loss of membrane integrity (Fig. 1a). Calcofluor White (CW) interacts with chitin leading to disruption of chitin polymer assembly and impairing the corresponding linkages to other cell wall components^35^. Increased susceptibility to Congo Red (CR) suggests a reduction of β-1,3 and β-1,6 glucans. Increased sensitivity to Sodium Dodecyl Sulphate (SDS) reflects a compromised membrane that may be due to differences in the chitin layer. Finally, we investigated whether loss of *A. fumigatus GNA1* enhanced the effects glucan synthesis inhibition by the echinocandin caspofungin (Fig. 1a). Exposure to caspofungin induced a terminal phenotype in the *Δgna1* mutant (*≥* 10^4^ conidia) and was fungicidal at lower concentrations. This provides evidence of synergistic *in vitro* activity of a beta glucan synthase inhibitor together with the *Δgna1* mutant.

To probe the effects of Glc on germination, parental and *Δgna1* mutant germlings were examined by light microscopy (Fig. 1c). Under conditions tested the *Δgna1* mutant was non-viable (Fig. 1a) and examination of the conidia demonstrated a subset that were enlarged and highly disorganised with evidence of multiple, rudimentary germ-tubes whose development was prematurely arrested (Fig. 1c). Despite macroscopically comparable phenotypes, the *Δgna1* mutant conidia incubated in the presence of 90 mM GlcNAc also displayed abnormalities compared to the parental strain (Fig. 1c). Although hyphae were evident at 24 h, there was also evidence of conidial enlargement and a lack of polarised hyphal growth with some *Δgna1* germ tubes unable to branch (Fig. 1c). We hypothesise that this sub-population of abnormal conidia accounts for the significant reduction in viability of the *A. fumigatus Δgna1* strain (Fig. 1c). GlcNAc is unable to completely rescue this phenotype. This suggests a possible cell wall phenotype due to reduced flux through the hexosamine biosynthetic pathway.

Defects in the cell surface architecture of the parental and *Δgna1* mutant strains were also revealed by both Scanning Electron Microscopy (SEM) and Transmission Electron Microscopy (TEM). SEM analysis demonstrated that the *Δgna1* mutant grown under terminal phenotype conditions (0.1 mM Glc and 1.5 μM GlcNAc) displayed conidiophore collapse (Fig. 1d). The vesicular head was grossly abnormal with the presence of “naked” phialides and no obvious conidia compared to the parental strain (Fig. 1d). TEM analysis revealed a disorganised cell wall ultrastructure for the *Δgna1* mutant, with a less defined chitin layer and a more dispersed outer mannoprotein layer (Fig. 1d). These data suggest that loss of *GNA1* is associated with a defective and potentially more porous cell wall and provides a molecular basis for the observed synergy with caspofungin. Taken together, these data show that the *Δgna1* strain displays a cell wall phenotype.

### GNA1 contributes to A. fumigatus virulence in models of aspergillosis

Ascertaining the physiological implications of *GNA1* loss by translating *in vitro* phenotypic characterisations into *in vivo* infection models is an important step in providing genetic validation as an antifungal target. In particular, it is critical to determine whether *in vivo* GlcNAc scavenging systems could compensate for *GNA1* loss. Non-vertebrate mini-host model systems such as the greater wax moth larvae (*Galleria mellonella*) represent a rapid initial screening strategy^36^. Larval inoculation resulted in a mean survival time of 3 days for the parental and reconstituted strains whereas the *Δgna1* mutant was avirulent with a survival profile indistinguishable from the PBS control group (Fig. 2a).

**Figure 2.**
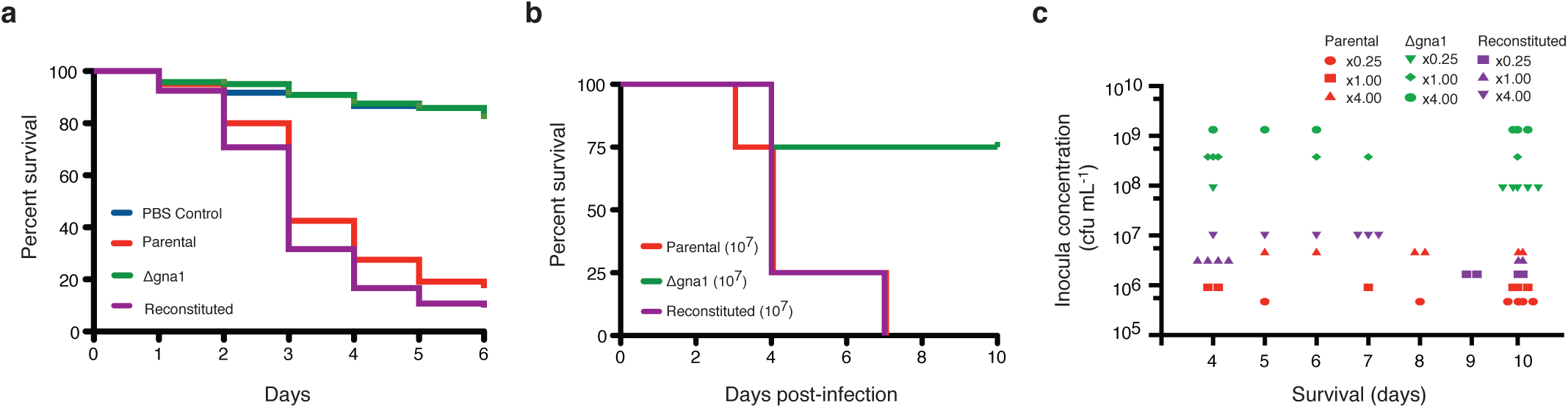
GNA1 contributes to A. fumigatus pathogenicity in invertebrate and murine models of aspergillosis. **a** *Galleria mellonella* were inoculated with 5 x 10^5^ conidia of *A. fumigatus* (parental, *Δgna1* mutant or reconstituted strains) or PBS as a vehicle control. Survival was monitored daily as evident by movement and lack of melanisation. Data is combined from four independent experiments (*n* = 4 with a combined total of 120 larvae per group). *P* < 0.0001. **b** Neutropenic male CD1 mice were exposed to 1 x 10^7^ *A. fumigatus* aerosolised conidia from the parental, *Δgna1* mutant and reconstituted strains in an inhalational chamber (*n* = 4 per group). **c** Neutropenic male CD1 mice were exposed to a trio of refined doses of *A. fumigatus* aerosolised conidia from the parental, *Δgna1* mutant and reconstituted strains in an inhalational chamber (*n* = 6 per group). A reference dose of x 1.00 was set to that observed to induce signs of disease from day 3. Each symbol represents an individual animal. In **b** and **c** mice were monitored for up to 10 days post infection and any mouse showing clinical signs of disease or distress was culled immediately and death being recorded as the next day.

A disadvantage of *G. mellonella* is that it fails to represent the natural inhalation route of *Aspergillus*. To recapitulate inhalational acquisition of *A. fumigatus* and provide insight into the physiological interplay of alveolar glucose and GlcNAc levels in circumventing *GNA1* loss, we used an established neutropenic murine model of invasive pulmonary aspergillosis^37^. A preliminary study was performed using three concentrations of viable *A. fumigatus* conidia and the contribution of the *Δgna1* mutant to pathogenicity was compared with the parental and reconstituted strains. In this experiment, the *Δgna1* mutant was less virulent (Fig. 2b). The median survival time was four days for the parental and reconstituted strains whereas a value could not be determined for the *Δgna1* mutant (75% survival). This finding was confirmed in a subsequent study (Fig. 2c) using doses refined for each strain designed to induce signs of invasive aspergillosis after approximately 3 days post inhalation (see Supplemental Experimental Methods). Although the mean survival times between *A. fumigatus* strains are not directly comparable due to differences in dose, there was an 83% survival rate to day 10 for mice exposed to the lowest concentration (x0.25) of the *Δgna1* mutant strain compared to 33% and 0% for the parental and reconstituted strains at their highest concentrations (x4.00, see Table S2). Approximately 1 to 2 log higher concentrations of *Δgna1* were required to induce clinical disease (Fig. 2c). Overall these findings suggest that a degree of metabolic (GlcNAc) rescue of *Δgna1* may occur *in vivo*. Nevertheless, these data show that *GNA1* contributes to *A. fumigatus* virulence in models of aspergillosis.

### Discovery of small molecules that target the Gna1 dimer interface

There are currently no selective small molecule inhibitors for the Gna1 class of enzymes to attempt to chemically phenocopy the genetic phenotype. Mechanistically, *A. fumigatus* (*Af*) Gna1 facilitates *N*-acetylation of GlcN-6P by promoting direct nucleophilic attack at the thioester carbonyl group of the acetyl co-enzyme A (AcCoA) cofactor by the sugar-phosphate amino group^38^. Although sharing only 30% sequence identity, structurally *Af*Gna1 and *Homo sapiens* (*Hs*) Gna1 adopt the same overall fold and domain architecture, with highly conserved regions localised to the AcCoA binding site with minor differences in the sugar substrate binding region. These differences are localised to amino acids involved in sugar-phosphate group recognition, which in the fungal enzyme utilise side chain functionality that is less polar in character^38,39^. In an attempt to circumvent issues of high structural homology between human and fungal Gna1, we deployed a fragment-based approach^40^ to discover small molecules that target previously unidentified pockets that are selective for the fungal enzyme. Bio-layer interferometry (BLI) was used to identify Gna1 binders from a 650 compound in-house fragment library^41–43^. This produced an initial hit rate of 5.7% (Figure S2). The top hit (**1**, Fig. 3a) displayed stoichiometric binding and high ligand efficiency (LE = 0.57 kcal mol^-1^ NHA^-1^ for **1**), with a 6 μM equilibrium dissociation constant which was confirmed by isothermal titration calorimetry (ITC) (Figure S2).

**Figure 3.**
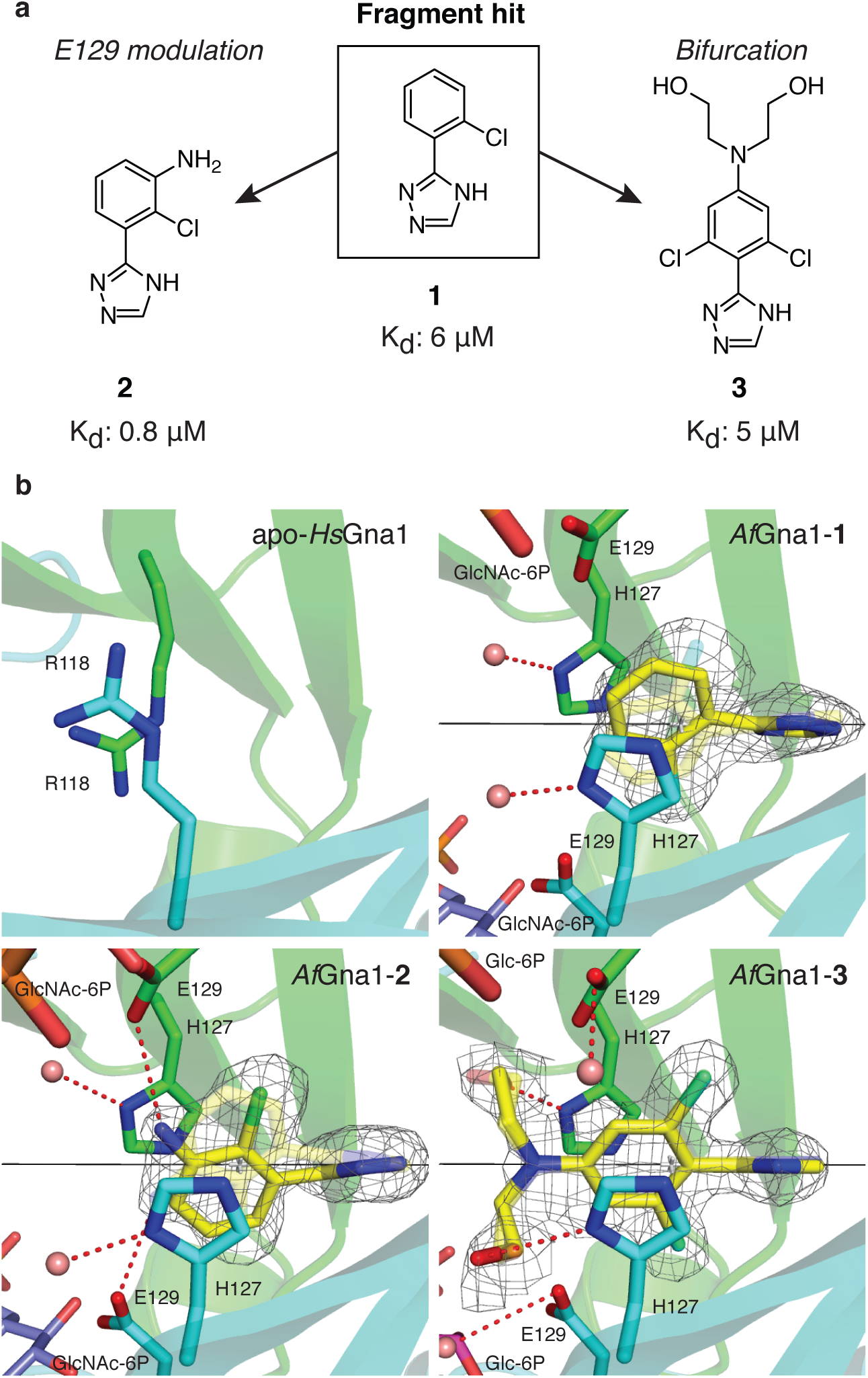
Derivatives of a fragment screen hit selectively target an unusual Gna1 pocket close to the active site. **a** Chemical structures and binding affinities of fragment-hit (**1**) and derivatives (**2** and **3**). **b** Panels showing the structures of apo-*Hs*Gna1 (PDB 2HUZ) and co-complex structures of *Af*Gna1-**1**, *Af*Gna1-**2** and *Af*Gna1-**3**. Gna1 monomer units (cyan and green); bound small molecules and crystallographic symmetry-related molecules (yellow and transparent sticks, respectively); GlcNAc-6P (blue sticks); Glc-6P (plum sticks); water molecules (pink spheres); Hydrogen-bond interactions (red dashed lines); crystallographic axis (black lines). In all co-complex structures, small molecule electron density was observed binding across the crystallographic two-fold axis that generates the *Af*Gna1 homodimer. The symmetry-equivalent molecules of fragment hit **1**, **2** and **3** are shown as transparent sticks. In apo-*Hs*Gna1, the novel dimer interface binding pocket is occluded by R118 residues preventing ligand binding. In *Af*Gna1, substitution of R118 by H127 facilitates π−π stacking with **1**, **2** and **3** forming the basis of fungal selective fragment binding. Superposition of enzymatic product, GlcNAc-6P (PDB 2VXK) onto co-complex structures *Af*Gna1-**1** and *Af*Gna1-**2** demonstrates the proximity of the fragment hit and derivatives to the E129 side chains and substrate binding sites. The AcCoA cofactor is not visible from selected viewpoint. The F_o_−F_c_ map for the bound small molecules (fragment hit **1**, **2** and **3**) are shown as a mesh (grey) contoured to 2.5 σ.

We next used X-ray crystallography to determine the binding mode of **1**. *Af*Gna1 was co-crystallised with AcCoA and soaked into solutions of **1**. Synchrotron diffraction data to 1.6 Å (Fig 3, Table 1) were collected and used to solve the structure by molecular replacement. Previous work has shown that *A. fumigatus* Gna1 possesses a classic GNAT α/β fold, forming an obligate homodimer. This dimer shows extensive secondary structure swapping with substrate and cofactor binding to the two active sites that are formed by channels on the surface of the enzyme, composed of amino acids from both *Af*Gna1 monomers^39^. Intriguingly, electron density corresponding to fragment hit **1** was observed in a pocket buried at the dimer interface in the core of the enzyme, binding across a crystallographic two-fold axis that generates the *Af*Gna1 homodimer and the symmetry-related molecule of fragment hit **1**, visible in the *Af*Gna1-**1** co-complex structure (Fig. 3b). This hydrophobic pocket (∼400 Å^3^) accomodates fragment hit **1** which is positioned ∼8 Å from both GlcN-6P substrate binding sites with a histidine residue (H127) from each monomer trapping the fragment in a π−π stacking arrangement (Fig. 3b). Critically, *Af*Gna1 H127 is substituted with an arginine (R118) in the human enzyme, blocking the pocket and importantly, preventing fragment binding to *Hs*Gna1 (Fig. 3b, Figure S3), whereas H127 is conserved amongst clinically relevant fungal species (Figure S3). With the exception of R118, the amino acids lining the fragment binding pocket are conserved between clinically relevant fungal Gna1 orthologues and *Hs*Gna1 (Figure S3).

**Table 1.** Crystallographic data collection and refinement statistics.

|  | *AfGna1-1-AcCoA* | *AfGna1-2-AcCoA* | *AfGna1-3-AcCoA-(Glc-6P)* |
| --- | --- | --- | --- |
| Beamline | *DLS-IO4-1* | *ESRF- ID30A-3* | *ESRF- ID30A-1* |
| Resolution (Å) | 56.46 (1.64 -1.58) | 40.32 (1.80 -1.74) | 58.2 (2.08 - 2.01) |
| Space group | C 2 2 21 | C 2 2 21 | C 2 2 21 |
| Unit cell (Å) *a, b, c* | 67.64, 102.51, 56.62 | 70.21, 101.2, 56.37 | 70.96, 101.71, 55.91 |
| $\alpha = \beta = \gamma$ (°) | 90 | 90 | 90 |
| Total reflections | 150350(5090) | 94486 (4877) | 53191 (2813) |
| Unique reflections | 26623 (2681) | 20528 (2069) | 13064 (1243) |
| Multiplicity | 5.6(3.8) | 4.6 (4.7) | 4.0 (4.2) |
| Completeness (%) | 97 | 98 | 94.6 |
| $I/\sigma(I)$ | 11.8(0.7) | 16.2 (2.3) | 14.5 (2.4) |
| $R_{\text{merge}}$ (%) | 6.2(123) | 4.1 (49.0) | 4.9 (57.8) |
| $CC_{1/2}$ | 99.8 | 99.9 | 99.9 |
| $R_{\text{work}}/R_{\text{free}}$ (%) | 20.0 / 22.7 | 19.0 / 23.5 | 21.1 / 26.4 |
| Average *B*-factor (Å<sup>2</sup>) | 38.0 | 35.7 | 46.6 |
| protein | 37.90 | 35.48 | 46.25 |
| ligand | 32.77 | 29.38 | 47.62 |
| solvent | 47.42 | 42.93 | 54.03 |
| No. of residues | 164 | 165 | 165 |
| No. of water mol. | 108 | 88 | 45 |
| *R.m.s deviations* |  |  |  |
| Bonds (Å) | 0.012 | 0.013 | 0.009 |
| Angles (°) | 1.89 | 1.89 | 1.81 |
| PDB code | 6TDH | 6TDG | 6TDF |

Statistics for the highest-resolution shell are shown in parentheses.

To develop the fragment scaffold towards *Af*Gna1-selective inhibition, the binding characteristics of the fragment within the dimer interface pocket were explored to identify the structural requirements for optimal *Af*Gna1 binding (Figure S2). The staggered bi-ring system of **1** in the *Af*Gna1-**1** complex is likely a consequence of reducing steric repulsion between the *ortho*-chloro substituent of the aryl ring system and the adjacent 1,2,4-triazole (dihedral angle, θ; ∼51°). This results in optimal positioning of **1** to facilitate the important π−π stacking interactions with the H127 residues present in the pocket (Fig. 3b). It appeared essential to maintain this bi-ring arrangement as loss or replacement of the *ortho*-substituent increased or ablated binding affinity (Figure S2).

With the *ortho*-chloro substituted scaffold identified we considered the close proximity of the glutamic acid residue (E129) that separates fragment hit **1** and the substrate binding site from each respective *Af*Gna1 monomer. Superposition of *Af*Gna1-GlcNAc-6P complex (the *N*-acetylated enzymatic product of GlcN-6P; PDB 2VXK), onto the co-complex structure of *Af*Gna1 with **1** illustrates the close proximity of the E129 side chains to the fragment hit **1** (3.4 Å) and to the substrate binding sites (8.2 Å) (Fig. 3b). Previous Gna1 studies have indicated that orthologous glutamate residues, whilst not playing a catalytic role, contribute to substrate and cofactor binding through direct contacts and through hydrogen-bond mediated water interactions^27,44,45^. Indeed, our data demonstrate that mutations which remove the E129 carboxylate (E129A) or substitute it with the corresponding amide (E129Q), reduced enzymatic activity compared to WT *Af*Gna1 (Figure S4). As such, we hypothesised that the close proximity of E129 residues to fragment hit **1** may offer scope to develop derivatives that exploit the E129 the side-chain functionality to both improve fragment binding affinity and elicit enzyme inhibition through perturbation of E129-mediated interactions (“E129 modulation approach”, Fig. 3a).

To explore the feasibility of harnessing E129-fragment proximity towards enzyme binding and inhibition, a *meta*-anilino group was installed into the scaffold. This resulted in fragment derivative **2**, which displayed sub-micromolar affinity (0.8 μM, Fig. 3a). Derivative **2** also displayed improved ligand efficiency compared to fragment hit **1** as a result of this minimal scaffold addition, resulting in a highly efficient binder (LE = 0.64 kcal mol^-1^ NHA^-1^ for **2**). Overall, **2** maintains the important π−π binding interaction observed in the *Af*Gna1-**1** complex structure. As observed in the *Af*Gna1-**2** complex structure, the introduction of the *meta*-anilino group causes a positional shift of **2** (32° from fragment hit **1**), facilitating a hydrogen bond interaction between **2** and the E129 side chain that likely drives the observed enhancement in binding affinity (Fig. 3). Although derivative **2** did not show inhibition against *Af*Gna1 in a recombinant enzyme assay, exploiting the proximity of the E129 side chains may be a route to improving binding potency whilst providing an anchoring point from which to introduce chemical functionality to modulate the position or protonation state of the E129 residues in order to achieve inhibition.

As an alternative approach to achieving *Af*Gna1 inhibition, we aimed to exploit the protein tunnels that connect the dimer interface fragment binding pocket to the substrate binding sites (distance of **1** to substrate binding site = ∼8 Å, Fig. 3). Extension of the fragment scaffold into the active sites by traversing these tunnels would effectively generate a competitive enzyme inhibitor by blocking substrate binding at both *Af*Gna1 active sites (“bifurcation approach”, Fig. 3a).

In contrast to E129 modulation, the success of bifurcation is predicated upon the ability of fragment hit **1** derivatives to both successfully reach the buried dimer interface binding pocket and be spatially accommodated despite the concomitant increase in fragment molecular size, which could conceivably hinder binding on both accounts. In order to test the ability of a bifurcated derivative of **1** to bind *Af*Gna1, **3** was synthesised. This bifurcated fragment introduces an *N,N*-diethyl alcohol moiety into the scaffold and displays a binding affinity of the same order as fragment hit **1** (5 μM, Fig. 3a, Figure S2). Importantly, this illustrates that the introduction of a bifurcated modification into the fragment scaffold is tolerated and that fragment binding affinity is maintained and not compromised by the introduction of simple linker groups. An *Af*Gna1-**3** complex structure, which additionally contains Glc-6P, indicates that **3** is accessible to the fragment binding site, despite narrow tunnels linking the fragment binding pocket to the enzyme active sites (Fig. 3b, Figure S3). Like fragment hit **1** and **2**, the important binding interactions and binding mode are maintained in the *Af*Gna1-**3**-(Glc-6P) complex structure. In addition, the hydroxy-functionality of the *N,N*-diethyl alcohol moiety displaces H127-coordinating water molecules and interacts with the acceptor nitrogen atoms of the H127 side chains (Fig. 3b). The introduction of a *para*-anilino anchoring point for bifurcation leverages the sp^2^ character of the anilino-nitrogen, maintaining the planarity necessary at the branch point for correct *N,N*-diethyl alcohol orientation whilst *ortho*-dichloro substitution positions the fragment along the 2-fold crystallographic axis (with a positional shift of 0.6 Å and 19° reorientation from fragment hit **1**). As a result, bifurcation vectors are identically orientated towards both *Af*Gna1 active sites in contrast to the binding positions of fragment hit **1** and **2** (Fig. 3b). Moreover, the positioning of the Glc-6P 6.2 Å from the ethyl chain linker of **3**, visible in the *Af*Gna1-**3**-(Glc-6P) complex structure (Fig. 3b, Figure S3), is suggestive of fragment-linking approaches towards *Af*Gna1 inhibition, albeit challenging due to the limited size of the tunnel^46^. In line with this, derivative **3** did not show inhibition against *Af*Gna1 in a recombinant enzyme assay, probably due to the inability of the *N,N*-diethyl alcohol moiety to extend into and block the enzyme active sites.

Globally, the structural differences between the co-complexes of *Af*Gna1 with **1**, **2** and **3** are negligible (RMSD ∼0.35 - 0.40). However, when considering buried protein pockets and cavities in general, such seemingly trivial displacements may be of significance. Recent molecular dynamics simulations and experiments have suggested that large scale side chain or secondary structure conformational changes are not a pre-requisite for ligand binding to buried cavities but rather subtle and rapid movement of peptide structural elements can be sufficient to facilitate ligand ingress and egress^47–50^. Although beyond the scope of this work, we speculate that the effects of any subtle movement of structural elements may propagate within *Af*Gna1 and be important for ligand binding dynamics to the unusual buried dimer interface pocket. Taken together, we have discovered small molecules that target a fungal specific binding hotspot at the Gna1 dimer interface, providing a scaffold for the development of Gna1 inhibitors.

## Discussion

The current clinical management of invasive fungal infections is multi-faceted and as a consequence of limited antifungal pipeline progression, clinicians must preserve the finite number of agents available to manage infection^51^. Recent decades witnessed an exodus of research and development into antifungal drug discovery by the pharmaceutical industry resulting in very few agents with novel mechanisms of action undergoing clinical trials. Clinical need prevails, despite optimal antifungal therapy, mortality rates remain around 50% although patient co-morbidities also contribute^1^. In response, a multidisciplinary approach to develop new targets has been advocated^52^.

Successful identification and validation of novel and ligandable targets is critical to the progression of the antifungal pipeline but the current shortage of unique targets combined with the intrinsic therapeutic hurdles introduced by the similarities of fungal and human enzymes only serves to complicate an already underappreciated and growing medical problem^1,2,53^. An attractive antifungal target is the cell wall, a structure unique to fungi and essential for survival. The sugar polymers constituting the cell wall are all synthesised from sugar nucleotide precursors. Here, we used a multidisciplinary approach focused on an essential step in the biosynthesis of the sugar nucleotide, UDP-GlcNAc, the precursor of cell wall chitin as a target against *A. fumigatus.* Cell wall chitin has long been proposed as an antifungal target. The nikkomycins and polyoxins are substrate analogues of UDP-GlcNAc and inhibit chitin synthesis *in vitro* with a predilection for *CHS-A* but fail *in vivo* primarily due to resistance in chitin synthase isoenzymes^54–56^. Our approach circumvents this failure by targeting Gna1, a GNAT-family enzyme with no isoforms. Inhibitors targeting other GNAT family members are known in the literature^57,58^, with bisubstrate inhibitors - molecules incorporating tethered mimics of both substrate and cofactor - offering improved selectivity profiles compared to alternative approaches but harbouring non-ideal pharmacokinetic properties^59–63^. However, such an approach is unlikely to offer robust selectivity in the case of Gna1 due to the high structural homology between fungal and human enzyme.

Previous failings in target-based drug discovery derive from an inadequate initial assessment of a molecular target in terms of physiological function^64^. Although *GNA1* was shown to be essential for *in vitro* growth in *S. cerevisiae* and *C. albicans*^27,29^, intrinsic differences in gene function across fungi due to diverging biological and physiological functions, genetic redundancies or scavenging pathways precludes predictions. Essentiality in yeasts does not automatically correlate in *A. fumigatus* and therefore this was experimentally validated to justify any potential investment in future inhibitor development. We demonstrated that genetic disruption of *A. fumigatus GNA1* results in a terminal phenotype under simulated physiological Glc and GlcNAc concentrations. This is further supported by our *in vivo* study using a neutropenic murine model of invasive aspergillosis with inhalation of *A. fumigatus Δgna1* approximately 100-fold less pathogenic than control strains. While the finding corroborates previous work (Mio et al., 2000), we speculate that earlier *in vitro* studies were not sufficiently robust to detect physiologically relevant nutritional rescue strategies since if a gene were genuinely “essential” it would result in avirulence rather than attenuation. Gene deletion strategies incorporating nutritional rescue do not permit different levels of *GNA1* expression to be directly investigated. Altering the levels of extracellular GlcNAc reflects the ability of *A. fumigatus* to induce salvage pathways.

Synergy between cell wall active agents is increasingly seen as an attractive option for antifungal treatment^65^. This is exemplified *in vitro* in *A. fumigatus* by the synergy between nikkomycin and the echinocandins^66^. At present the echinocandins are advocated as salvage therapy in invasive aspergillosis in part due to their fungistatic activity^67^. We show that *A. fumigatus* Δ*gna1* combined with caspofungin has a fungicidal effect on *in vitro* growth completely abolishing not only the terminal phenotype but also growth under maximal GlcNAc rescue. This provides further evidence that combining a β-glucan synthase inhibitor together with chitin inhibition is synergistic *in vitro*. A *GNA1* inhibitor could enhance the efficacy of the echinocandins and offer additional treatment modalities potentially as a resistance sparing agent. In terms of biological importance, Gna1 is an attractive antifungal target.

In addition to our genetic validation of Gna1, we sought to identify compounds that would chemically phenocopy the biological phenotype. The inherent difficulty in leveraging inhibitor potency and selectivity between high-homology enzyme families amongst higher eukaryotic organisms is a challenging problem to fully resolve. This applies to Gna1 where deletion in *A. fumigatus* yields a terminal phenotype, but deletion of the murine orthologue leads to developmental phenotypes^68^. We pursued a fragment-based approach to overcome this challenge by discovering a previously unidentified fungal selective pocket within *Af*Gna1, positioned on the two-fold axis generating the Gna1 biological homodimer. Crucially, this pocket is absent in the human orthologue, where it is occupied by larger side chains (arginines, versus histidines in the fungal enzyme).

Our data show that fragment derivatives can recapitulate binding to this unusual fragment anchoring site and rational design targeting this pocket allows the development of micromolar binders that do not target the human enzyme. Although not yet displaying *Af*Gna1 inhibitory activity at high micromolar concentrations, the opportunities towards fungal-selective *Af*Gna1 inhibition opened up by the discovery of fragments binding at this fungal-selective binding pocket is promising.

Moreover, with the exception of R118, the amino acids that constitute the interface binding pocket are well conserved between clinically relevant fungal Gna1 orthologues and *Hs*Gna1. Although sequence alignments have limited predictive capacity in determining the structural context of the interface pocket and its ligandability, this high degree of conservation warrants further structural and chemical exploration. This binding pocket may offer a ligandable handle against other fungal Gna1 orthologues, in particular, *C. auris* which recently became the first fungal species listed as an urgent drug resistant threat^11^.

Identifying and pursuing novel avenues for inhibitor development to overcome organism selectivity barriers is critical to reversing the dearth in the antifungal pipeline as is the timely identification and validation of unusual and novel targets, exemplified herein by Gna1.

The work reported here also serves to further exemplify the ability of fragment-based screening to uncover unique and otherwise unforeseen avenues towards enzyme inhibition through the discovery of novel binding sites. Successful fragment campaigns that provide the foundation to potent inhibitors commonly begin with fragments that display no intrinsic enzymatic inhibitory activity^69^. The fact that fragment hit **1** and derivatives (**2** and **3**) display a stable binding mode is often an important pre-requisite to subsequent successful fragment development campaigns^70^. Moreover, while small molecule therapeutics that bind to protein interfaces are uncommon, such approaches have been successfully utilised in structure-based design campaigns to inhibit enzyme targets. For example, HIV-1 and HIV-2 protease inhibitors, such as Darunavir, bind to a cavity at the protein dimer interface located adjacent to the enzyme active site, forming contacts to the interface cavity and inhibiting enzymatic activity through hydrogen-bonding to catalytically important D25 carboxylate side-chains of both protease monomers thereby competitively inhibiting viral polypeptide access^71–73^.

The *Af*Gna1-**3**-(Glc-6P) complex structure offers a glimpse at potential avenues of inhibition via a combination of bifurcation and fragment-linking approaches. Although a rare and challenging form of fragment development, successful fragment-linking campaigns ideally utilise the conjugation of fragment pairs with contrasting hydrophobic and polar ligand-protein binding characteristics^46^. Such characteristics are exemplified herein with fragment hit **1** and the sugar (or pseudo-sugar) substrate and therefore provide an incentive to further explore this approach. In parallel, exploring potential routes towards E129 modulation of substrate binding has led to the synthesis of a sub-micromolar fragment binder and a key chemical scaffold which can now be further explored and modified to elicit the desired inhibitory response.

In summary, we have provided genetic validation of *Af*Gna1 as an antifungal target and have discovered small molecules that target the Gna1 dimer interface, providing a platform for generating inhibitors of the enzyme. As demonstrated, our multidisciplinary approach, incorporating both genetic and chemical target assessment, is critical in determining target feasibility. Such approaches are not limited to fungal pathogens and are applicable to other infectious agents and are of particular relevance when considering targets with high sequence and structural homology to human enzymes.

## Acknowledgements

We wish to thank the Dundee Drug Discovery Unit for access to the Fragment library and the European Synchrotron Radiation Facility, Grenoble and Diamond Light Source, Oxford for time at the beamline. The assistance from Mr Martin Kierans, School of Life Sciences, University of Dundee with the Electron Microscopy is gratefully acknowledged. This work was supported by a Wellcome Trust Postdoctoral Research Training Fellowship for Clinicians (WT105772/A/14/Z) to DL and an MRC Programme Grant (MR/M004139/1) to DMFvA. DB was funded by a University of Aberdeen Summer Research Scholarship. The structures have been deposited in the Protein Data Bank with accession codes 6TDH, 6TDG and 6TDF.

## Author Contributions

DMFvA, DEA, MS conceived the project. DEA, MS and DMFvA designed the experiments. DEA and MS executed the experiments with assistance from OGR, DAR, DB, DRS, ATF and WF. DEA, MS and DMFvA interpreted the data. DEA, MS and DMFvA wrote the manuscript with input from other authors.

## Materials and methods

### Ethics statement

All animal experiments were performed by Evotec Ltd (UK) under UK Home Office Licence 40/3644 and ethically approved by The University of Manchester Standing Committee.

### Strains, culture medium and solutions

The genetic lineage and nutritional growth requirements of all strains used and generated in this work is provided in Table S1. Further details are provided in the Supplemental Experimental Methods.

### Generation of A. fumigatus Δgna1 mutant

To delete *GNA1*, construct p*Δgna1pyrG*+ was designed to replace the entire 644 bp coding region of *GNA1* with the URA-blaster (also called the *pyrG* blaster) as a reusable selection marker by homologous recombination^74^ (Figure S1). Refer to the Supplemental Experimental Methods for all PCR primer sequences used in this work. The Supplemental Experimental Methods contain details of the experimental process including protoplast transformation, phenotypic screening, PCR screening and Southern Blot verification (Figure S1) of *A. fumigatus Δgna1*.

### Generation of an A. fumigatus GNA1+6H::pyrG+ reconstituted strain

A construct p*GNA1*+6H was designed to replace the entire URA-blaster (8.3 kb) in the Δ*gna1* mutant with the native *GNA1* sequence by homologous recombination (Figure S1). Experimental procedures culminating in Southern Blot verification followed those for generation of the Δ*gna1* mutant (see Table S1 and Supplemental Experimental Methods). To permit *in vivo* applications by restoring uridine and uracil autotrophy, the reconstituted strain (*GNA1*+6H::*pyrG*-) was complemented with *pyrG* by random integration. Primers were used to amplify a 1.9 kb portion of the *A. fumigatus pyrG* gene^75^ from pXDRFP4^76^. Refer to Supplemental Experimental Methods for further details.

### In vitro phenotypic analysis of A. fumigatus Δgna1 mutant

An agar dilution method was used to investigate *in vitro* growth in the presence of different concentrations of glucose and GlcNAc. Complete medium (CM) containing 90 mM, 150 μM and 1.5 μM GlcNAc was supplemented with 0.1 mM Glucose (Glc). CM supplemented with *≥* 50 mM Glc and 0 mM GlcNAc was a Δ*gna1* negative growth control. For preliminary evaluation of growth characteristics under the above conditions, suspensions containing serial dilutions of conidia (5 μl, 1 x 10^6^ - 1 x 10^3^) from the respective *A. fumigatus* strains were inoculated on solid medium and incubated at 37 **°**C for 48 h.

To further assess the interaction between Glc and GlcNAc on growth, an agar microdilution method^77^ was modified using flat-bottomed 96-well plates to generate a checkerboard assay. This enabled simultaneous testing of 285 different combinations of Glc and GlcNAc (final concentration range 0 mM to 50 mM, 2-fold serial dilutions) in a 200 μl final volume (180 μl of non-glucose CM agar and 10 μl each of Glc and GlcNAc). Each well represented a different growth condition and was inoculated with *A. fumigatus* viable conidia (1 x 10^3^) and incubated at 37 **°**C for 48 h. Wells were scored independently by two unblinded investigators using a numerical code (1, no growth/inoculum spotting to 7, growth equivalent to positive control) to obtain an average score per condition. Scores were converted to colours generating a growth heatmap (Excel, Microsoft).

Germination was examined by inoculating liquid medium (20 ml, CM 110 mM Glc and CM 0.1 mM Glc + 90 mM GlcNAc) with conidia (1 x 10^5^) and incubating in a Petri dish containing glass coverslips. At specified time-points (5 h, 8 h, 12 h and 24 h), the coverslips were removed, washed and fixed (2 ml, 3.7 % paraformaldehyde, 20 min). Coverslips were mounted onto glass slides and examined using a light microscope (Leica ICC50 HD, ×40 objective lens magnification).

To examine the response of the *A. fumigatus Δgna1* mutant to cell wall perturbing agents, suspensions of conidiophores (5 μl, 1 x 10^6^ - 1 x 10^2^) were inoculated onto solid 0.1 mM Glc CM + *≥* 50 mM GlcNAc in the presence of Calcofluor White, Congo Red, Sodium Dodecyl Sulphate (SDS) (25 - 100 μg/ml) and Caspofungin (0.0625 - 0.5 μg/ml). Following incubation (48 h), plates were examined. The experiment was performed three times.

For analysis of cell wall architecture, conidia and mycelia were grown on solid medium, fixed, processed and examined using transmission and scanning electron microscopy. Refer to Supplemental Experimental Methods for further details.

### A. fumigatus Δgna1 infection models

Greater wax moth larvae, *Galleria mellonella*, (Livefoods Direct, Sheffield, UK) were used as an infection model^36,78,79^. Larvae were maintained at room temperature in wood shavings in the dark^80^ and those in the 6^th^ instar measuring 2.0 cm in length (approximately 250 mg) were randomly assigned into groups. The *A. fumigatus* parental, Δ*gna1* and reconstituted strains were inoculated into the hind proleg of larvae (*n* = 30 per group) as a 10 μl PBS suspension of 5 x 10^5^ resting conidia. Negative controls were inoculated with PBS only. Larvae were protected from light, incubated at 37 °C and examined at 24 h intervals. Mortality, defined by lack of movement in response to stimuli and discolouration (melanisation) of the cuticle, was scored for 6 days.

An inhalational neutropenic murine model of invasive aspergillosis^37,81^ was performed by Evotec Ltd (UK) (see ethics statement). As this was an initial proof of concept study and due to insufficient published literature using the parental strain in this particular murine model, group size by power analyses were unable to be performed for statistical differences between strains. Sharing the data generated in this study will inform future power calculations.

Male CD1 mice (Charles River, UK) weighing 23-26 g were used for all experiments and allowed to acclimatise for at least 7 days. Mice were randomly assigned to groups and housed in sterilised individually ventilated cages with HEPA filtered sterile air and sterile aspen chip bedding (changed every 3-4 days). Food and water were provided *ad libitum*. The room temperature was 22 °C (±1 °C) with a relative humidity of 60% and maximum background noise of 56 dB. Mice were exposed to 12 h light/dark cycles.

Mice were rendered neutropenic with intraperitoneal cyclophosphamide (250 mg/kg on day -2 relative to infection, and 200 mg/kg on day +3) and sub-cutaneous cortisone acetate (200 mg/kg on day -2 and day +3). To generate conidial suspensions of the target inoculum (Table S2) for inhalational inoculation (infection) of mice, *A. fumigatus* parental, Δ*gna1* and reconstituted strains were prepared according to a National Institute of Health (NIH) standard operating procedure^82^ with nutritional requirements as per Table S1. Viable cell counts were performed on serially diluted aliquots of each conidial suspension.

Each cage was randomly assigned to a study group and infected by an inhalation chamber system^37^. On the day of infection, mice (*n* = 4 per group for experiment 1; *n* = 6 per group for experiment 2) were exposed for 1 h to 12 ml of PBS containing *A. fumigatus* conidia aerosolised via a Micro Mist® nebuliser (Teleflex Medical, Kernen, Germany) in an acrylic chamber. Enrofloxacin (50 ppm, 0.2 ml/water bag) was administered from day -3 to day +10 to prevent bacterial infection. Mice were weighed and monitored at least daily and the study refined using surrogate markers for invasive pulmonary aspergillosis. Any mouse with signs of illness and distress* were culled immediately (terminal anaesthesia or cervical dislocation) and death recorded as being the next day.

* A scoring system was used based on the following parameters: weight loss; hypothermia less than 32 °C; appearance (hunching, poor coat condition); clinical signs: respiration (noisy laboured breathing); tremors/convulsions; unprovoked behaviour (little peer interaction) and provoked behaviour (subdued when stimulated).

### Statistical Analyses

Biological experiments were performed at least three independent times with the exception of electron microscopy. Statistical analyses were carried out using GraphPad Prism (v5.0b). Data is expressed as the mean ± SD (Fig. 1c). To compare more than two groups, one-way ANOVA test with Bonferroni’s multiple comparison test was used (Fig 1c). For comparison of survival curves, the Kaplan-Meier Log-rank (Mantel-Cox) test was performed (Fig. 2a only).

### AfGna1 protein expression and purification

*Af*Gna1 was purified as described earlier^39^. Briefly, the sequence encoding the protein was inserted into a pGEX6P-1 vector for expression of a GST tagged fusion that contains PreScission protease cleavage site. For protein expression, the plasmid was transformed into *Escherichia coli* BL21(DE3) pLysS. Cell cultures were grown to an OD600 of 0.8 and expression was initiated by the addition of 250 μM IPTG at 17 °C. The cultures were then allowed to grow for a further 16 h before harvesting by centrifugation. After lysis, affinity chromatography and GST tag cleavage, eluted protein was concentrated and further purified by size exclusion chromatography using a Superdex 75, 26/60 column. Corresponding fractions confirmed by SDS-PAGE were pooled concentrated, flash frozen in liquid nitrogen and stored at −80 °C.

### AfGna1 fragment screen and binding affinity measurements

To identify a potential fragment binder of *Af*Gna1, the fragment library of the University of Dundee Drug Discovery Unit (DDU) comprising of 652 structurally diverse compounds was screened using biolayer interferometry (BLI)^41–43^. BLI experiments were performed on an Octet Red 384 system (Forte Bio, USA) using super streptavidin (SSA) biosensors and biotinyated-*Af*Gna1 Primary hits identified from initial screening were defined as fragments with a response rate greater than three robust standard deviation units above the median. Confirmation of fragment hit binding was performed with a follow-up concentration dose-response series. Binding isotherms were fitted and visualised using Octet software (Forte Bio, USA) to derive dissociation constants (K_d_). Refer to supporting information for further experimental details.

### AfGna1 crystallography

*Af*Gna1 crystallization was carried out as described earlier^39^. Crystallisation was manually performed using a sitting drop approach by combining protein solution previously incubated with AcCoA for 2 h on ice with reservoir solution in a 1:1 ratio and equilibrated against 60 μl reservoir solution. *Af*Gna1 crystals resembling bipyramidal or bar shaped were generally observed after 72 h incubation at 20 °C.

For fragment soaking, a soaking solution for each confirmed fragment hit was prepared by combining a fragment DMSO stock solution (50-200 mM) with crystallisation reservoir solution. After incubation (1 hr – overnight) at 20 °C, crystals were transferred to a cryoprotectant using a nylon loop, immersed for 10 s and flash frozen in liquid nitrogen.

Diffraction data were collected at the European Synchrotron Radiation Facility (ESRF), Grenoble (France) and the Diamond Light Source, Oxford (UK). The diffraction data were automatically processed and scaled by *xia*2^83^. An initial *Af*Gna1-**1** complex was solved by molecular replacement using *MOLREP*^84^ using *Af*Gna1 apo structure (PDB ID 2VEZ) as a search molecule. Refinement was done using REFMAC5^85^ and model building with Coot^86^. Well-defined electron density for fragment hit **1** was evident at the dimer interface. An apo version of this new model served as a search molecule for the determination of subsequent complexes by molecular replacement. Where appropriate, ARP/wARP^87^ was used to partially build the model before further refinement. PRODRG^88^ was used to generate coordinates and topologies of the compounds (ligands). Ligands were not included until their conformations were completely defined by unbiased σA-weighted |*F*_o_|-|*F*_c_|, ϕ_calc_ electron density maps.

### Chemical synthesis of fragment derivatives

#### 2-chloro-3-(1,2,4-triazol-3-yl)aniline, Compound **2**

3-amino-2-benzoic acid (0.5 g, 2.90 mmol) was dissolved in dichloromethane/*N,N*-dimethylformamide (7 ml:2 ml). To the solution was added *N,N*-Diisopropylethylamine (1.5 ml, 8.70 mmol), 1-Hydroxybenzotriazole hydrate (470 mg, 3.48 mmol) and *N*-(3-Dimethylaminopropyl)-*N-*ethylcarbodiimide hydrochloride (667 mg, 3.48 mmol). The solution was stirred at room temperature under argon over 20 min, ensuring that all solid material had dissolved. Ammonium hydroxide solution (25 %, 0.35 ml) was added dropwise to the stirring solution, forming a pale-yellow suspension which was then left overnight to stir. After aqueous work-up (∼5-10 ml ddH_2_O), extraction with Ethyl acetate (4 x 15 ml) and drying over sodium sulphate, the crude material (∼440 mg) was concentrated and rapidly purified over a silica plug (Ethyl acetate → Ethyl acetate/MeOH). Eluted fractions were analysed by LCMS, confirming product formation and fractions containing the desired intermediate amide product were combined (320 mg, 64% yield). Without further purification, the intermediate amide (314 mg, 1.84 mmol) was dissolved in *N,N*-Dimethylformamide dimethyl acetal (6 ml) and was refluxed, under argon at 115 °C for 15 h. After refluxing, the solution was cooled and evaporated to dryness *in vacuo*. In a separate receptacle, 50-60 % Hydrazine hydrate solution (150 μL) was added slowly (dropwise) to acetic acid (9 ml). This solution was then rapidly transferred to the dried intermediate and refluxed at 95 °C for 7 hours. After reflux, the solution was cooled and evaporated to dryness *in vacuo*. The crude material was dry-loaded onto silica and was directly purified by Flash Chromatography (heptane/ethyl acetate → ethyl acetate → ethyl acetate/MeOH). Eluted fractions were analysed by LCMS, appropriately combined, evaporated to dryness and resulted in an off-white solid (post-lyophilisation from 50 % acetonitrile and ddH_2_O) (52 mg, 14% yield, Rf 0.76, ethyl acetate). Overall, 9% yield over 2 steps.

^1^H NMR (500 MHz, MeOH-D4): *δ =* 8.57 (s, 1 H), 7.21 (dd, J = 7.6 Hz, 7.6 Hz, 1 H), 7.06 (dd, J = 8.1 Hz, 1.6 Hz, 1H), 7.01 (dd, J = 7.5 Hz, 1.6 Hz, 1H).

^13^C NMR (101 MHz, MeOH-D4): *δ* = 156.64, 147.33, 145.86, 128.61, 125.90, 121.43, 119.22, 118.60.

HRMS-TOF: Theo. m/z 195.0432; obs. m/z 195.0428; [M+H], C_8_H_8_ClN_4_.

#### 2,2’-((3,5-dichloro-4-(1,2,4-triazol-3-yl)phenyl)azanediyl)bis(ethan-1-ol), Compound **3**

2,6-dichloro-4-nitrobenzamide (1 g, 4.25 mmol) was dissolved in ethanol (12 ml) followed by the addition of tin (II) chloride dihydrate (4.8 mg, 21.30 mmol). The resulting solution was refluxed at 70°C overnight at which point the solution was cooled, poured into ice water (50 ml) and pH adjusted (to pH ∼8-9) using 4N aqueous sodium hydroxide, where it was allowed to stir for 30 min. The resulting solid was removed by filtering over celite and the aqueous filtrate was then extensively extracted with ethyl acetate (6 x 25 ml). The organic phases were combined, dried over sodium sulphate and evaporated to dryness *in vacuo*, resulting in an off-white solid (intermediate aniline) which was used without further purification (72% yield).

The intermediate aniline (250 mg, 1.22 mmol) was dissolved in acetic acid (2.5 ml), followed by the addition of ∼3 M ethylene oxide-THF solution (2.5 ml). The resulting solution was stirred at room temperature for 24 h at which point the solvent was evaporated to near dryness *in vacuuo.* The crude material was dry-loaded onto silica and was directly purified by Flash Chromatography (ethyl acetate → ethyl acetate/MeOH). Eluted fractions were analysed by LCMS, appropriately combined and evaporated to dryness, resulting in a mixture of unreacted starting material, *N-* and *N,N-*ethyl alcohol products. The reaction and subsequent FLASH purification were then repeated using the combined unreacted starting material and mono-functionalised product to improve the conversion to the *N,N-*ethyl alcohol product (145 mg, 40% yield).

The *N,N-*ethyl alcohol product (100 mg, 0.34 mmol) was dissolved in a 1:1 mixture of pyridine and acetic anhydride (10 ml) and was left to stir overnight at which point the solution was evaporated to dryness followed by trituration of the resulting oil with toluene (2 x 5 ml) (∼quant. hydroxyl group protection). Without further purification, the *O*-acetyl protected material (∼125 mg, ∼0.34 mmol) was dissolved in *N,N*-dimethylformamide dimethyl acetal (5 ml) and was refluxed, under argon at 115 °C for 24 h. After refluxing, the solution was cooled and evaporated to dryness *in vacuo*. In a separate receptacle, 50-60 % hydrazine hydrate solution (8 eq., 85 μL) was added slowly (dropwise) to acetic acid (8 ml). This solution was then rapidly transferred to the dried intermediate and was refluxed at 95 °C for 15 hours. After reflux, the solution was cooled and evaporated to dryness *in vacuo*. The crude material was dry-loaded onto silica and was directly purified by Flash Chromatography (heptane/ethyl acetate → ethyl acetate). Eluted fractions were analysed by LCMS and appropriately combined and evaporated to dryness, resulting in a white foam (85 mg, 48% yield). The resulting 1,2,4-triazole (40 mg, 0.10 mmol) was then *O*-acetyl deprotected using a 1:1 solution of 1,4-dioxane and 1.5 M aqueous sodium hydroxide (pH ∼10). The reaction was monitored by TLC and on completion, the basic solution was neutralised using Amberlite IR-120 H^+^ ion-exchange resin. The resin was rinsed and the resulting solution was lyophilised. The lyophilised material was then purified by Flash Chromatography (2 g silica; gradient elution, 1:1 ethyl acetate/Heptane → ethyl acetate → 3:1 ethyl acetate/Methanol). Eluted fractions were analysed by LCMS and appropriately combined and evaporated to dryness (20 mg, 63% yield; Rf 0.41, ethyl acetate), with an overall yield of 9% yield over 5 steps.

^1^H NMR (500 MHz, MeOH-D4): *δ* = 8.34 (br, s, 1 H), 6.88 (s, 2H), 3.76 (t, J = 6.0 Hz, 4 H), 3.58 (t, J = 6.0 Hz, 4 H).

^13^C NMR (101 MHz, MeOH-D4): *δ* = 153.21-151.14 (br, s), 144.66, 137.53, 133.52, 112.14, 59.95, 54.58.

HRMS-TOF: Theo. m/z 317.0567; obs. m/z 317.0573; [M+H], C_12_H_15_Cl_2_N_4_O_2_.

## References

1. Brown, G. D. et al. Hidden killers: Human fungal infections. Sci. Transl. Med. 4, 165rv13 (2012).

2. Fisher, M. C., Hawkins, N. J., Sanglard, D. & Gurr, S. J. Worldwide emergence of resistance to antifungal drugs challenges human health and food security. Science (80-.). 360, 739–742 (2018).

3. Walsh, T. J. et al. Treatment of Aspergillosis: Clinical Practice Guidelines of the Infectious Diseases Society of America. Clin. Infect. Dis. 46, 327–360 (2008).

4. Agarwal, R. et al. Allergic bronchopulmonary aspergillosis: Review of literature and proposal of new diagnostic and classification criteria. Clin. Exp. Allergy 43, 850–873 (2013).

5. Denning, D. W. Global fungal Burden. Mycoses 56, 13 (2013).

6. Schauwvlieghe, A. F. A. D. et al. Invasive aspergillosis in patients admitted to the intensive care unit with severe influenza: a retrospective cohort study. Lancet Respir. Med. 6, 782–792 (2018).

7. Hamill, R. J. Amphotericin B formulations: A comparative review of efficacy and toxicity. Drugs 73, 919–934 (2013).

8. Allen, D., Wilson, D., Drew, R. & Perfect, J. Azole antifungals: 35 years of invasive fungal infection management. Expert Rev. Anti. Infect. Ther. 13, 787–798 (2015).

9. Holt, S. L. & Drew, R. H. Echinocandins: Addressing outstanding questions surrounding treatment of invasive fungal infections. Am. J. Heal. Pharm. 68, 1207–1220 (2011).

10. Vermes, A. Flucytosine: a review of its pharmacology, clinical indications, pharmacokinetics, toxicity and drug interactions. J. Antimicrob. Chemother. 46, 171–179 (2000).

11. CDC. Antibiotic Resistance Threats in the United States, 2019. www.cdc.gov/DrugResistance/Biggest-Threats.html (2019).

12. Chowdhary, A., Kathuria, S., Xu, J. & Meis, J. F. Emergence of Azole-Resistant Aspergillus fumigatus Strains due to Agricultural Azole Use Creates an Increasing Threat to Human Health. PLoS Pathog. 9, (2013).

13. Head, M. G., Fitchett, J. R., Atun, R. & May, R. C. Systematic analysis of funding awarded for mycology research to institutions in the UK, 1997-2010. BMJ Open 4, e004129 (2014).

14. Roemer, T. & Krysan, D. J. Antifungal drug development: challenges, unmet clinical needs, and new approaches. Cold Spring Harb. Perspect. Med. 4, a019703 (2014).

15. Perfect, J. R. The antifungal pipeline: A reality check. Nat. Rev. Drug Discov. 16, 603–616 (2017).

16. Falci, D. R. & Pasqualotto, A. C. Profile of isavuconazole and its potential in the treatment of severe invasive fungal infections. Infect. Drug Resist. 6, 163–174 (2013).

17. Han, D. H. Isavuconazole Noninferior to Voriconazole in Invasive Mold Disease. Monthly prescribing reference October 12th (2014).

18. Buil, J. B. et al. Isavuconazole susceptibility of clinical Aspergillus fumigatus isolates and feasibility of isavuconazole dose escalation to treat isolates with elevated MICs. J. Antimicrob. Chemother. 73, 134–142 (2018).

19. Gastebois, A., Clavaud, C., Aimanianda, V. & Latgé, J. P. Aspergillus fumigatus: Cell wall polysaccharides, their biosynthesis and organization. Future Microbiol. 4, 583–595 (2009).

20. Tada, R., Latge, J.-P. & Aimanianda, V. Undressing the Fungal Cell Wall/Cell Membrane - the Antifungal Drug Targets. Curr. Pharm. Des. 19, 3738–3747 (2013).

21. Vetting, M. W. et al. Structure and functions of the GNAT superfamily of acetyltransferases. Arch. Biochem. Biophys. 433, 212–226 (2005).

22. Selitrennikoff, C. P., Allin, D. & Sonneborn, D. R. Chitin biosynthesis during Blastocladiella zoospore germination: evidence that the hexosamine biosynthetic pathway is post translationally activated during cell differentiation. Proc. Natl. Acad. Sci. U. S. A. 73, 534–538 (1976).

23. Milewski, S., Gabriel, I. & Olchowy, J. Enzymes of UDP-GlcNAc biosynthesis in yeast. Yeast 23, 1–14 (2006).

24. Fang, W. et al. Genetic and structural validation of Aspergillus fumigatus N-acetylphosphoglucosamine mutase as an antifungal target. Biosci. Rep. 33, 689–699 (2013).

25. Fang, W. et al. Genetic and structural validation of Aspergillus fumigatus UDP-N-acetylglucosamine pyrophosphorylase as an antifungal target. Mol. Microbiol. 89, 479– 493 (2013).

26. Hu, W. et al. Essential gene identification and drug target prioritization in Aspergillus fumigatus. PLoS Pathog. 3, e24 (2007).

27. Mio, T., Yamada-Okabe, T., Arisawa, M. & Yamada-Okabe, H. Saccharomyces cerevisiae GNA1, an essential gene encoding a novel acetyltransferase involved in UDP-N-acetylglucosamine synthesis. J. Biol. Chem. 274, 424–429 (1999).

28. Becker, J. M. et al. Pathway analysis of Candida albicans survival and virulence determinants in a murine infection model. Proc. Natl. Acad. Sci. U. S. A. 107, 22044–22049 (2010).

29. Mio, T., Kokado, M., Arisawa, M. & Yamada-Okabe, H. Reduced virulence of Candida albicans mutants lacking the GNA1 gene encoding glucosamine-6-phosphate acetyltransferase. Microbiology 146, 1753–1758 (2000).

30. Boehmelt, G. et al. Cloning and characterization of the murine glucosamine-6-phosphate acetyltransferase EMeg32. Differential expression and intracellular membrane association. J. Biol. Chem. 275, 12821–12832 (2000).

31. Cova, M. et al. The Apicomplexa-specific glucosamine-6-phosphate N-acetyltransferase gene family encodes a key enzyme for glycoconjugate synthesis with potential as therapeutic target. Sci. Rep. 8, (2018).

32. D’Enfert, C. et al. Attenuated virulence of uridine-uracil auxotrophs of Aspergillus fumigatus. Infect. Immun. 64, 4401–4405 (1996).

33. Williams, V. & del Poeta, M. Role of Glucose in the Expression of Cryptococcus neoformans Antiphagocytic Protein 1, App1. Eukaryot. Cell 10, 293–301 (2011).

34. Jain, R. et al. The MAP kinase MpkA controls cell wall integrity, oxidative stress response, gliotoxin production and iron adaptation in Aspergillus fumigatus. Mol. Microbiol. 82, 39– 53 (2011).

35. Ram, A. F. J. et al. The cell wall stress response in Aspergillus niger involves increased expression of the glutamine: Fructose-6-phosphate amidotransferase-encoding gene (gfaA) and increased deposition of chitin in the cell wall. Microbiology 150, 3315–3326 (2004).

36. Slater, J. L., Gregson, L., Denning, D. W. & Warn, P. A. Pathogenicity of Aspergillus fumigatus mutants assessed in Galleria mellonella matches that in mice. Med. Mycol. 49, (2011).

37. Sheppard, D. C. et al. Novel Inhalational Murine Model of Invasive Pulmonary Aspergillosis. Antimicrob. Agents Chemother. 48, 1908–1911 (2004).

38. Hurtado-Guerrero, R., Raimi, O., Shepherd, S. & van Aalten, D. M. F. Glucose-6-phosphate as a probe for the glucosamine-6-phosphate N-acetyltransferase Michaelis complex. FEBS Lett. 581, 5597–5600 (2007).

39. Hurtado-Guerrero, R. et al. Structural and kinetic differences between human and Aspergillus fumigatus D-glucosamine-6-phosphate N-acetyltransferase. Biochem. J. 415, 217–223 (2008).

40. Erlanson, D. A., Fesik, S. W., Hubbard, R. E., Jahnke, W. & Jhoti, H. Twenty years on: The impact of fragments on drug discovery. Nature Reviews Drug Discovery vol. 15 605–619 (2016).

41. Wartchow, C. A. et al. Biosensor-based small molecule fragment screening with biolayer interferometry. J. Comput. Aided. Mol. Des. 25, 669–676 (2011).

42. Shah, N. B. & Duncan, T. M. Bio-layer interferometry for measuring kinetics of protein-protein interactions and allosteric ligand effects. J. Vis. Exp. e51383 (2014).

43. Prakash, O. et al. Identification of Leishmania major UDP-Sugar Pyrophosphorylase Inhibitors Using Biosensor-Based Small Molecule Fragment Library Screening. Molecules 24, 996 (2019).

44. Peneff, C., Mengin-Lecreulx, D. & Bourne, Y. The Crystal Structures of Apo and Complexed Saccharomyces cerevisiae GNA1 Shed Light on the Catalytic Mechanism of an Amino-sugar N-Acetyltransferase. J. Biol. Chem. 276, 16328–16334 (2001).

45. Wang, J., Liu, X., Liang, Y. H., Li, L. F. & Su, X. D. Acceptor substrate binding revealed by crystal structure of human glucosamine-6-phosphate N-acetyltransferase 1. FEBS Lett. 582, 2973–2978 (2008).

46. Ichihara, O., Barker, J., Law, R. J. & Whittaker, M. Compound design by fragment-linking. Mol. Inform. 30, 298–306 (2011).

47. Feher, V. A., Baldwin, E. P. & Dahlquist, F. W. Access of ligands to cavities within the core of a protein is rapid. Nat. Struct. Biol. 3, 516–521 (1996).

48. Wang, Y., Papaleo, E. & Lindorff-Larsen, K. Mapping transiently formed and sparsely populated conformations on a complex energy landscape. Elife 5, (2016).

49. Lopata, A. et al. A hidden active site in the potential drug target mycobacterium tuberculosis dUTPase is accessible through small amplitude protein conformational changes. J. Biol. Chem. 291, 26320–26331 (2016).

50. Mondal, J., Ahalawat, N., Pandit, S., Kay, L. E. & Vallurupalli, P. Atomic resolution mechanism of ligand binding to a solvent inaccessible cavity in T4 lysozyme. PLoS Comput. Biol. 14, (2018).

51. Haiduven, D. Nosocomial aspergillosis and building construction. Med. Mycol. 47, S210–216 (2009).

52. Steinbach, W. J. Are We There Yet? Recent Progress in the Molecular Diagnosis and Novel Antifungal Targeting of Aspergillus fumigatus and Invasive Aspergillosis. PLoS Pathog. 9, (2013).

53. Perlin, D. S., Rautemaa-Richardson, R. & Alastruey-Izquierdo, A. The global problem of antifungal resistance: prevalence, mechanisms, and management. The Lancet Infectious Diseases vol. 17 e383–e392 (2017).

54. Gaughran, J. P., Lai, M. H., Kirsch, D. R. & Silverman, S. J. Nikkomycin Z is a specific inhibitor of Saccharomyces cerevisiae chitin synthase isozyme Chs3 in vitro and in vivo. J. Bacteriol. 176, 5857–5860 (1994).

55. Zhang, D. & Miller, M. J. Polyoxins and nikkomycins: progress in synthetic and biological studies. Curr. Pharm. Des. 5, 73–99 (1999).

56. Lenardon, M. D., Munro, C. A. & Gow, N. A. R. Chitin synthesis and fungal pathogenesis. Curr. Opin. Microbiol. 13, 416–423 (2010).

57. Huang, M., Huang, J., Zheng, Y. & Sun, Q. Histone acetyltransferase inhibitors: An overview in synthesis, structure-activity relationship and molecular mechanism. Eur. J. Med. Chem. 178, 259–286 (2019).

58. Wapenaar, H. & Dekker, F. J. Histone acetyltransferases: challenges in targeting bi-substrate enzymes. Clin. Epigenetics 8, (2016).

59. Lau, O. D. et al. HATs off: Selective synthetic inhibitors of the histone acetyltransferases p300 and PCAF. Mol. Cell 5, 589–595 (2000).

60. Zheng, Y. et al. Selective HAT Inhibitors as Mechanistic Tools for Protein Acetylation. Methods Enzymol. 376, 188–199 (2003).

61. Gao, F. et al. Synthesis and structure-activity relationships of truncated bisubstrate inhibitors of aminoglycoside 6′-N-acetyltransferases. J. Med. Chem. 49, 5273–5281 (2006).

62. Gao, C. et al. Rational design and validation of a Tip60 histone acetyltransferase inhibitor. Sci. Rep. 4, (2014).

63. Ngo, L., Brown, T. & Zheng, Y. G. Bisubstrate inhibitors to target histone acetyltransferase 1. Chem. Biol. Drug Des. 93, 865–873 (2019).

64. G. Wyatt, P., H. Gilbert, I., D. Read, K. & H. Fairlamb, A. Target Validation: Linking Target and Chemical Properties to Desired Product Profile. Curr. Top. Med. Chem. 11, 1275– 1283 (2011).

65. Campitelli, M., Zeineddine, N., Samaha, G. & Maslak, S. Combination Antifungal Therapy: A Review of Current Data. J. Clin. Med. Res. 9, 451–456 (2017).

66. Verwer, P. E. B., Van Duijn, M. L., Tavakol, M., Bakker-Woudenberg, I. A. J. M. & Van De Sande, W. W. J. Reshuffling of Aspergillus fumigatus cell wall components chitin and β-glucan under the influence of caspofungin or nikkomycin Z alone or in combination. Antimicrob. Agents Chemother. 56, 1595–1598 (2012).

67. Patterson, T. F. et al. Practice Guidelines for the Diagnosis and Management of Aspergillosis: 2016 Update by the Infectious Diseases Society of America. Clin. Infect. Dis. 63, e1–e60 (2016).

68. Boehmelt, G. et al. Decreased UDP-GIcNAc levels abrogate proliferation control in EMeg32-deficient cells. EMBO J. 19, 5092–5104 (2000).

69. Scott, D. E., Coyne, A. G., Hudson, S. A. & Abell, C. Fragment-based approaches in drug discovery and chemical biology. Biochemistry 51, 4990–5003 (2012).

70. Drwal, M. N. et al. Structural Insights on Fragment Binding Mode Conservation. J. Med. Chem. 61, 5963–5973 (2018).

71. Cardinale, D. et al. Homodimeric Enzymes as Drug Targets. Curr. Med. Chem. 17, 826– 846 (2010).

72. Da Silva, F., Bret, G., Teixeira, L., Gonzalez, C. F. & Rognan, D. Exhaustive Repertoire of Druggable Cavities at Protein–Protein Interfaces of Known Three-Dimensional Structure. J. Med. Chem. 62, 9732–9742 (2019).

73. Kovalevsky, A. Y., Louis, J. M., Aniana, A., Ghosh, A. K. & Weber, I. T. Structural Evidence for Effectiveness of Darunavir and Two Related Antiviral Inhibitors against HIV-2 Protease. J. Mol. Biol. 384, 178–192 (2008).

74. D’Enfert, C. Selection of multiple disruption events in Aspergillus fumigatus using the orotidine-5’-decarboxylase gene, pyrG, as a unique transformation marker. Curr. Genet. 30, 76–82 (1996).

75. Jiang, H., Shen, Y., Liu, W. & Lu, L. Deletion of the putative stretch-activated ion channel Mid1 is hypervirulent in Aspergillus fumigatus. Fungal Genet. Biol. 62, 62–70 (2014).

76. Yang, L. et al. Rapid production of gene replacement constructs and generation of a green fluorescent protein-tagged centromeric marker in Aspergillus nidulans. Eukaryot. Cell 3, 1359–1362 (2004).

77. Golus, J., Sawicki, R., Widelski, J. & Ginalska, G. The agar microdilution method – a new method for antimicrobial susceptibility testing for essential oils and plant extracts. J. Appl. Microbiol. 121, 1291–1299 (2016).

78. Renwick, J., Daly, P., Reeves, E. P. & Kavanagh, K. Susceptibility of larvae of Galleria mellonella to infection by Aspergillus fumigatus is dependent upon stage of conidial germination. Mycopathologia 161, 377–384 (2006).

79. Fallon, J. P., Troy, N. & Kavanagh, K. Pre-exposure of Galleria mellonella larvae to different doses of Aspergillus fumigatus conidia causes differential activation of cellular and humoral immune responses. Virulence 2, 413–421 (2011).

80. Mowlds, P. & Kavanagh, K. Effect of pre-incubation temperature on susceptibility of Galleria mellonella larvae to infection by Candida albicans. Mycopathologia 165, 5–12 (2008).

81. Sheppard, D. C. et al. Standardization of an experimental murine model of invasive pulmonary aspergillosis. Antimicrob. Agents Chemother. 50, 3501–3503 (2006).

82. NIH-NIAID-N01-AI-30041. Standard Operating Procedure for Murine Inhalational Pulmonary Aspergillosis: New Animal Models for Invasive Aspergillosis (IA). Version 1.10.

83. Winter, G. Xia2: An expert system for macromolecular crystallography data reduction. J. Appl. Crystallogr. 43, 186–190 (2010).

84. Vagin, A. & Teplyakov, A. MOLREP: An Automated Program for Molecular Replacement. J. Appl. Crystallogr. 30, 1022–1025 (1997).

85. Vagin, A. A. et al. REFMAC5 dictionary: Organization of prior chemical knowledge and guidelines for its use. Acta Crystallogr. Sect. D Biol. Crystallogr. 60, 2184–2195 (2004).

86. Emsley, P., Lohkamp, B., Scott, W. G. & Cowtan, K. Features and development of Coot. Acta Crystallogr. Sect. D Biol. Crystallogr. 66, 486–501 (2010).

87. Langer, G., Cohen, S. X., Lamzin, V. S. & Perrakis, A. Automated macromolecular model building for X-ray crystallography using ARP/wARP version 7. Nat. Protoc. 3, 1171–1179 (2008).

88. Schüttelkopf, A. W. & Van Aalten, D. M. F. PRODRG: A tool for high-throughput crystallography of protein-ligand complexes. Acta Crystallogr. Sect. D Biol. Crystallogr. 60, 1355–1363 (2004).

