## Supplemental Information for "Targeting a critical step in fungal hexosamine biosynthesis"

### Supplemental Figures

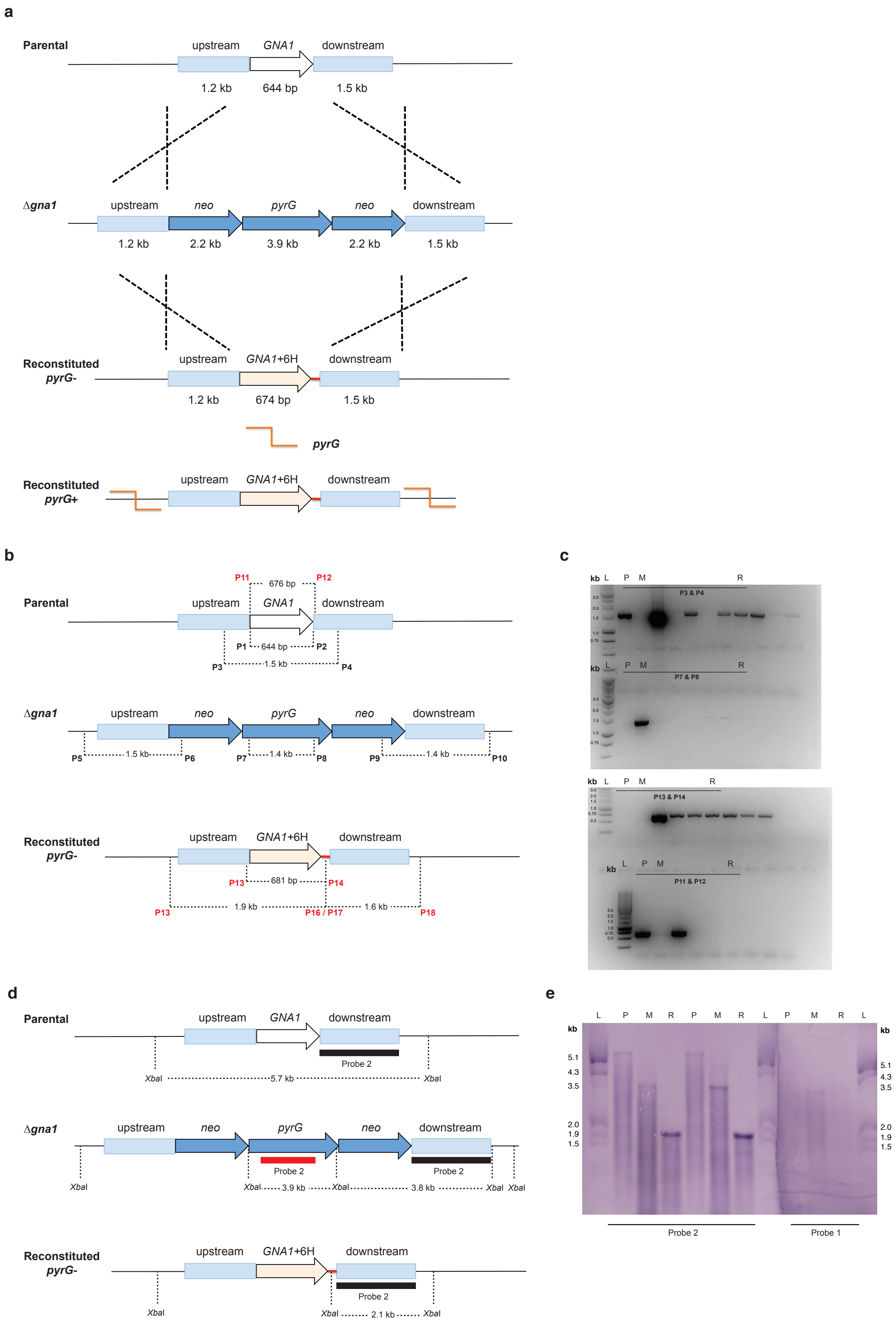

**Figure S1**. Refer to Figure 1.

**a.** ***A. fumigatus*** **gene deletion and reintegration design strategies to generate the strains for this study (see Supplemental Table 1).** Upper panels, *A. fumigatus* GNA1 gene deletion to generate a Δgna1 by homologous recombination. The GNA1 open reading frame was replaced with the URA blaster containing a pyrG selection marker. Middle panels, reintegration of GNA1 by homologous recombination generating an auxotrophic (pyrG-) reconstituted strain. The introduction of an engineered linker (red line) containing a polyhistidine tag (6H) and XbaI restriction site permits differentiation from the parental strain during PCR screening and Southern Blot analysis. Lower panels, reintegration of pyrG into the reconstituted strain by random chromosomal integration of the *A. fumigatus* pyrG gene. Black dashed lines illustrate the upstream and downstream flanking regions for homologous recombination (double crossover).

**b. PCR screening of transformants.** Upper panel, location of primer pairs for PCR with expected product sizes. Primer numbers correspond to those listed in the *Supplemental experimental methods*. Primers were designed to (i) span the junctions of homologous recombination; (iii) demonstrate replacement of *GNA1* with the URA blaster containing the *pyrG* selection marker and (iv) reintegration of *GNA1*+6H to differentiate between the *A. fumigatus* parental, Δ*gna1* and reconstituted strains.

**c. Representative raw data scans of agarose gel electrophoresis using four primer pairs during PCR screening of reconstituted strain transformants**. All primer pairs were used for screening. L, molecular weight DNA marker; P, parental, M, Δ*gna1* mutant; R, reconstituted (*pyrG-*). Unlabelled lanes represent transformants that failed at least one PCR screen.

**d. Southern Blot verification of** ***A. fumigatus*** **Δ*gna1*** **and reconstituted strain (*pyrG*** **deficient).** Upper panel, hybridisation probes were designed to differentiate between the *A. fumigatus* parental, Δ*gna1* and reconstituted strain (*pyrG-*). The *Xba*I cleavage sites are highlighted together with the expected size following probe hybridisation.

**e. Southern Blot membrane.** The membrane was cut during the experiment for each probe to hybridise (left, probe 2; right probe 1). L, DIG labelled DNA molecular weight marker III (Roche); P, parental; M, Δ*gna1* mutant; R, reconstituted (*pyrG-*) *A. fumigatus* strains.

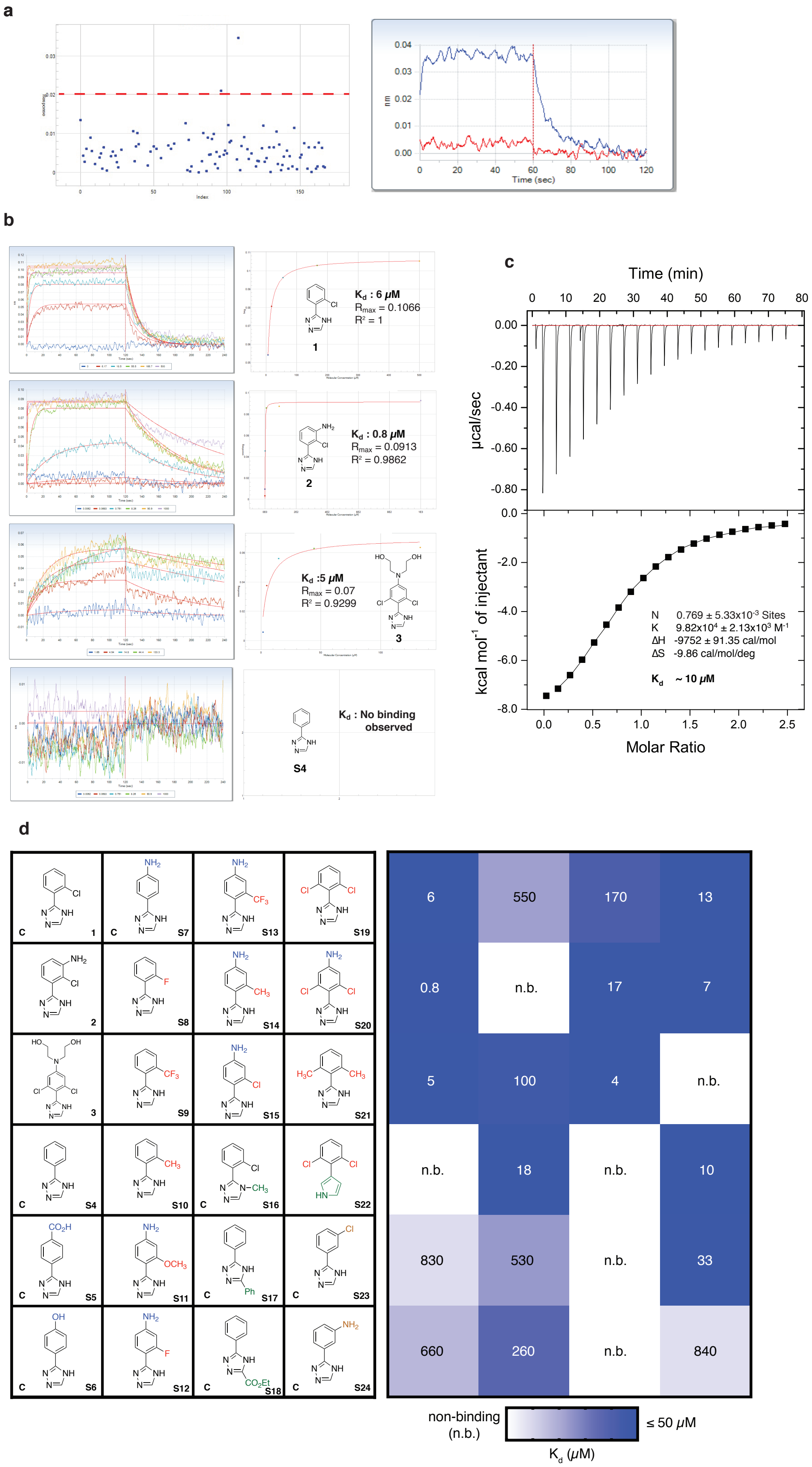
a
Index
0.04
0.03
0.02
0.01
0.00
nm
Time (sec)
b
$K_d$ : 6 $\mu$M
$R_{max}$ = 0.1066
$R^2$ = 1
1
$K_d$ : 0.8 $\mu$M
$R_{max}$ = 0.0913
$R^2$ = 0.9862
2
$K_d$ :5 $\mu$M
$R_{max}$ = 0.07
$R^2$ = 0.9299
3
$K_d$ : No binding observed
S4
c
Time (min)
µcal/sec
kcal mol$^{-1}$ of injectant
Molar Ratio
N 0.769 ± 5.33x10$^{-3}$ Sites
K 9.82x10$^4$ ± 2.13x10$^3$ M$^{-1}$
ΔH -9752 ± 91.35 cal/mol
ΔS -9.86 cal/mol/deg
$K_d$ ~ 10 $\mu$M
d
1
S7
S13
S19
2
S8
S14
S20
3
S9
S15
S21
S4
S10
S16
S22
S5
S11
S17
S23
S6
S12
S18
S24
6
550
170
13
0.8
n.b.
17
7
5
100
4
n.b.
n.b.
18
n.b.
10
830
530
n.b.
33
660
260
n.b.
840
non-binding (n.b.)
≤ 50 $\mu$M
$K_d$ ($\mu$M)

**Figure S2.** Refer to figure 3.

**a. Representative BLI responses observed for a subset of Drug Discovery Unit fragment library compounds (blue squares) assayed against *Af*Gna1.** Left panel, preliminary hits above the response rate cut-off of > 0.02 nm (red dashed line) were followed up. Initial hit rate of 5.7 % (37/650 at 200 µM [fragment]). Right panel, binding isotherms for preliminary hit, **1**; association (0 - 60 s) and dissociation (61 - 120 s) between immobilised *Af*Gna1 and fragment (blue line) or control (red line). Binding isotherm produced by fragment hit **1** was selected for further analysis based on examination of the fragment response rate difference between the association and dissociation curve.

**b. Representative BLI dose-response traces for fragment hit 1 and derivatives 2, 3 and S4.** Kinetic binding isotherms (left panels; concentration series [µM] vs. response (nm)) and binding analysis (right panels). Fragment hit **1** (upper), **2** (upper middle), **3** (lower middle) display binding behaviour towards biotinylated-*Af*Gna1. No binding was detected for commercial derivative **S4** using BLI analysis.

**c. Representative ITC binding isotherm confirming interaction between fragment hit 1 and *Af*Gna1**. The observed binding affinity $K_d$ ($1/K = K_d$) is of the same order of magnitude as $K_d$ derived from BLI binding analysis ($K_d$ : 6 µM and 10 µM from BLI and ITC, respectively).

**d. Binding analysis of fragment hit 1 derivatives towards *Af*Gna1**. $K_d$ was determined from BLI analysis. Compound-specific $K_d$ values are highlighted within the heatmap (right) and correspond to fragment hit **1** derivatives (left). Compounds are classed as binding to *Af*Gna1 (blue) or displaying non-detectable binding (white, n.b.). Compounds were synthesised using known literature procedures unless denoted by '**C**' (commercially available). The absence of *ortho*-substitution generally increases $K_d$ or ablates binding (**S4** – **S7**, **S23**, **S24**). Increasing the steric bulk of the *ortho*-substituent increases $K_d$ and abrogates binding (**S8** – **S10**). Introduction of a *para*-anilino group is tolerated (**S1**, **S15** vs. **S19**, **S20**) but does not compensate for steric bulk of *ortho*-substituents (**S11** – **S15**, **S21**). *Ortho*-chloro substitution (**1**, **2**, **3**, **S15**, **S19**, **S20**) corresponds to higher affinity binding interactions compared to structurally analogous derivatives (**S8** – **S14**, **S21**). Substitution of the 1,2,4-triazole ring ablates binding affinity (**S16** – **S18**).

**a**

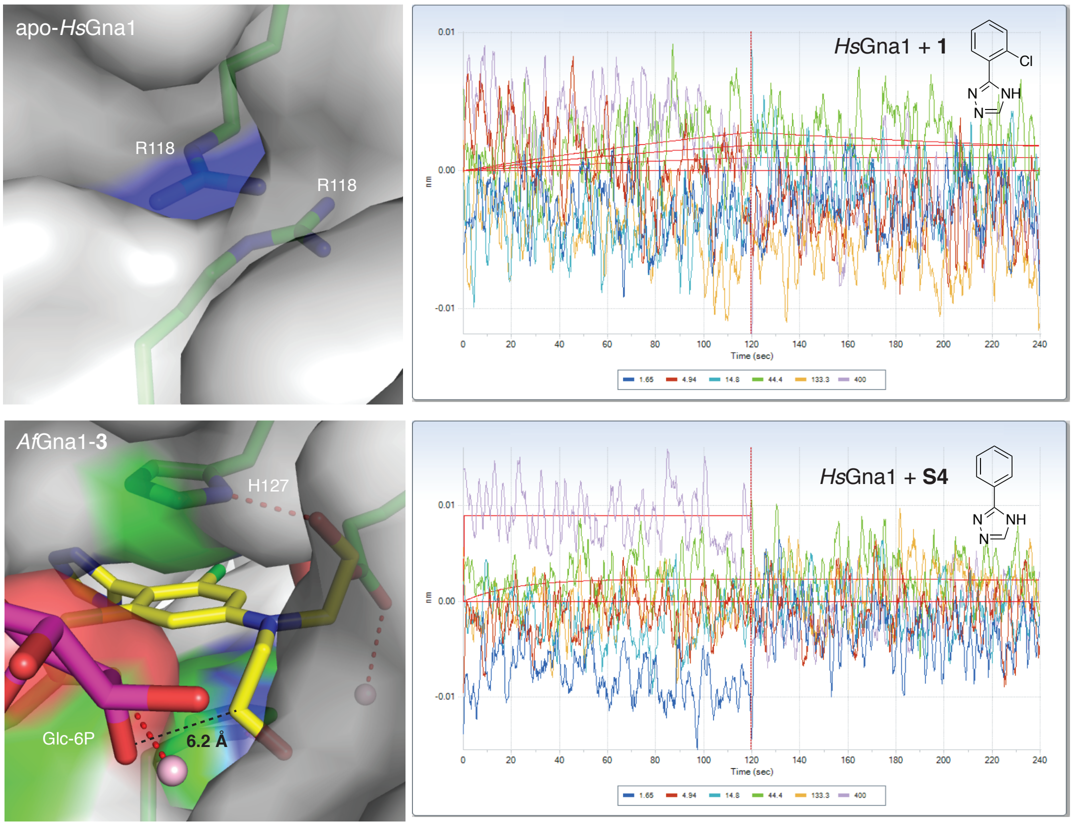


**b**

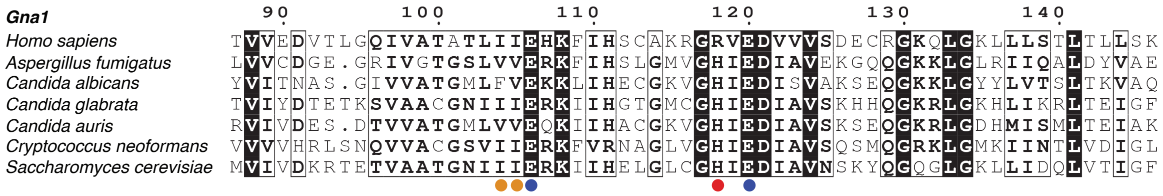


**Figure S3.** Refer to figure 3.

**a. Fragment hit 1 does not bind to *Hs*Gna1.** Repesentative BLI dose-response binding isotherms (upper right, lower right) of biotinylated-*Hs*Gna1 with fragment hit **1** (upper right) and **S4** (lower right). No binding is detected between fragment hit **1** and *Hs*Gna1 indicating the fungal selective nature of fragment binding as observed by the lack of dose-dependent binding isotherms. As with *Af*Gna1, **S4** is also non-binding to *Hs*Gna1. Surface representations of Apo-*HsGna1* (upper left) and the co-complex structure, *Af*Gna1-**3** (lower left). Gna1 monomers (grey surface, green sticks), derivative **3** (yellow sticks) and psuedo-substrate, Glc-6P (plum sticks). The dimer interface binding pocket is occulded in *Hs*Gna1 due to presence of the R118 residues blocking the pocket (upper left). In *Af*Gna1, R118 is substituted by H127 and the resulting fragment binding pocket is accessible by narrow tunnels from the substrate binding site (lower left). Derivative **3** is clearly visible in *Af*Gna1-**3**, with Glc-6P situated 6.2 Å from the *N*-ethyl alcohol moiety (lower left).

**b. Sequence alignment of *Hs*Gna1, *Af*Gna1 and other clinically relevant fungal species demonstrating a near conserved binding pocket.** Sequence alignment reveals the *Af*Gna1 H127 residue, important for the π-π stacking interaction with **1**, is conserved across the fungal species indicated but is absent in *Hs*Gna1 (red circle), where it is substituted to arginine (R118). Amino acids of the interface binding pocket are well conserved across other clinically relevant fungal species (blue circle = identical; orange circle = similar). Sequence alignment was generated using ENDscript[1].

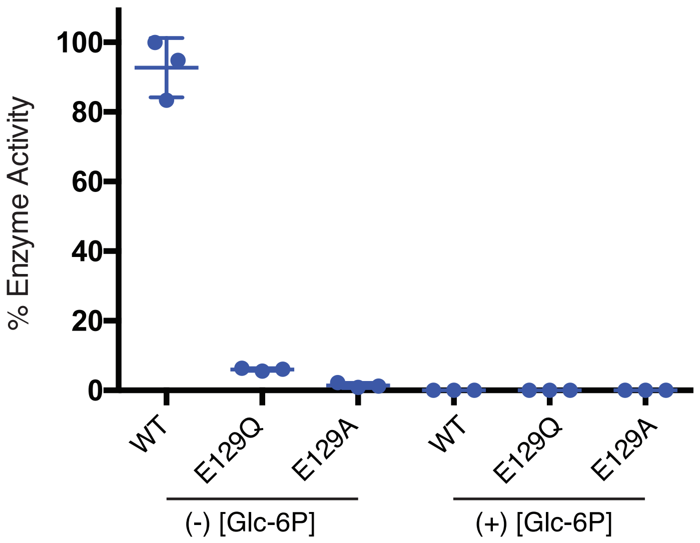


**Figure S4. E129 mutation reduces *Af*Gna1 enzymatic activity.** Refer to Figure 3.
The glutamate residue corresponding to *Af*Gna1 E129 in Gna1 orthologues has been shown to play important roles in substrate and cofactor binding. The introduction of mutations into *Af*Gna1 that remove the negative charge of the E129 carboxylate group (E129A) or substitute it with the corresponding amide (E129Q) reduce *Af*Gna1 enzymatic activity to ≲10% of wild-type (WT) levels. This suggests that perturbation of the E129 residues is a valid strategy to elicit enzymatic inhibition through modulation of substrate (and possibly cofactor) interactions.
Enzyme activity was assessed by a fluorometric thiol-detection assay as previously reported[2]. Enzymatic turnover releases CoA which then reacts with fluorogenic dye, 7-Diethylamino-3-(4-maleimidophenyl)-4-methylcoumarin (CPM, Sigma-Aldrich) allowing quantification of enzyme activity (ex/em; 384/470 nm).
Each data point represents % enzyme activity. Activity of the mutant enzymes are indicated as a percentage of WT *Af*Gna1. Error bars represent the standard deviation of replicate measurements ($n$ = 1). Pseudo-substrate, Glc-6P (100 mM) was used as a control to confirm enzyme inhibitory capacity. The WT and the mutant *Af*Gna1 proteins (E129A, E129Q) were expressed as a GST-fusion, purified with Glutathione-Sepharose beads followed by GST-tag cleavage (precision protease) according to published procedures.

#### Supplemental Tables

**Table S1.** ***A. fumigatus*** **strains used and generated in this work.** Related to Figure 1.
Strains were inoculated on minimal medium (MM) or complete medium (CM) with modifications to the base media as indicated (+, addition; -, removal). Any variation in glucose and *N*-acetylglucosamine (GlcNAc) concentrations in the growth culture medium for phenotypic analysis is indicated in the *Supplemental Experimental Methods*.

| Strain | Genotype and description | Specific nutritional requirements* | Reference |
|---|---|---|---|
| *Ku80*Δ | CEA17*akuB*$^{KU80}$*::PYRG+*<br>**Parental strain for *in vitro* and *in vivo* experiments.** | Nil | da Silva Ferreira et al., (2006) |
| *Ku80Δpyr G-* (Auxotrop hic) | CEA17*akuB*$^{KU80}$::PYRG+ PYRG-<br>**Parental recipient strain for protoplast transformation.** | + 5 mM uridine & 5 mM uracil | da Silva Ferreira et al., (2006) |
| Δ*gna1* | CEA17*akuB*$^{KU80}$::PYRG+ PYRG- ::*gna1* PYRG+<br>**Gene deletion, knockout, mutant.** | + ≥ 50 mM GlcNAc<br>- Glc | This study. |
| *Ku80*Δ6H *pyrG-* *GNA1*+6H (Auxotrop hic) | CEA17*akuB*$^{KU80}$::PYRG+ PYRG- ::*gna1*+6H<br>**Reconstituted strain derived from Δ*gna1* for Southern Blot.** | + 5 mM uridine & 5 mM uracil | This study. |
| *Ku80Δ6H pyrG+* *GNA1*+6H | CEA17*akuB*$^{KU80}$::PYRG+ PYRG- ::*gna1*+6H ::*Af*PYRG+<br>**Reconstituted strain for *in vitro* and *in vivo* phenotypic analysis.**<br>**Derived from *Ku80*Δ6H*pyrG*-.** | Nil | This study. |
| YJ-407 | China General Microbiological Culture Collection Centre 0386<br>**Wild-type strain for Southern Blot.** | Nil | Xia et al., (2001) |

**Table S2. Study groups for proof of concept of pathogenicity of *A. fumigatus* Δ*gna1* in a murine neutropenic model of inhalational invasive aspergillosis.** Related to Figure 2. Experiment 1 ($n$ = 4 per group), preliminary study to investigate the pathogenicity of each *A. fumigatus* strain at three different target concentrations (10-fold difference). Experiment 2 ($n$ = 6 per group), refined study using a narrower range of concentrations (based on the results of experiment 1) to investigate pathogenicity. Doses (a), (b) and (c) represent x0.25, x1.00 and x4.00 of an estimated dose inducing signs of disease from day 3 post infection.

| Experiment (group) | *A. fumigatus* strain | Target concentration (cfu $mL^{-1}$) | Inoculum viable count (cfu $mL^{-1}$) |
|---|---|---|---|
| 1 (1a) | Parental | $1.0 \times 10^6$ | $1.45 \times 10^6$ |
| 1 (1b) | Parental | $1.0 \times 10^7$ | $1.29 \times 10^7$ |
| 1 (1c) | Parental | $1.0 \times 10^8$ | $1.32 \times 10^8$ |
| 1 (2a) | Δ*gna1* | $1.0 \times 10^6$ | $7.55 \times 10^5$ |
| 1 (2b) | Δ*gna1* | $1.0 \times 10^7$ | $7.95 \times 10^6$ |
| 1 (2c) | Δ*gna1* | $1.0 \times 10^8$ | $7.75 \times 10^7$ |
| 1 (3a) | Reconstituted | $1.0 \times 10^6$ | $1.10 \times 10^6$ |
| 1 (3b) | Reconstituted | $1.0 \times 10^7$ | $1.17 \times 10^7$ |
| 1 (3c) | Reconstituted | $1.0 \times 10^8$ | $1.09 \times 10^8$ |
| 2 (1a) | Parental | $2.5 \times 10^5$ | $4.7 \times 10^5$ |
| 2 (1b) | Parental | $1.0 \times 10^6$ | $9.1 \times 10^5$ |
| 2 (1c) | Parental | $4.0 \times 10^6$ | $4.8 \times 10^6$ |
| 2 (2a) | Δ*gna1* | $7.5 \times 10^7$ | $8.9 \times 10^7$ |
| 2 (2b) | Δ*gna1* | $3.0\ 10^8$ | $3.8 \times 10^8$ |
| 2 (2c) | Δ*gna1* | $1.2 \times 10^9$ | $1.3 \times 10^9$ |
| 2 (3a) | Reconstituted | $1.2 \times 10^6$ | $1.7 \times 10^7$ |
| 2 (3b) | Reconstituted | $5.0 \times 10^6$ | $3.2 \times 10^6$ |
| 2 (3c) | Reconstituted | $2.0 \times 10^7$ | $1.0 \times 10^7$ |

#### Supplemental Experimental Methods

##### Strains, culture medium and solutions

*A. fumigatus* strains were maintained as 20% glycerol stocks at -80 °C (long term) or conidial suspensions in sterile water at 4 °C (short term). For strain propagation approximately 20 µl of *A. fumigatus* conidia and 100 µl of sterile water were inoculated onto solid medium (Table S1) and incubated at 37 °C for at least 72 h until sporulation. Conidia were harvested using a sterile spreader by flooding the surface of each plate with 10 ml of 0.01% Tween 20 in phosphate buffered saline (PBST). After recovery into a sterile tube, the suspension was centrifuged at 4000 rpm for 10 min followed by removal of the supernatant; the pellet was washed twice in sterile water. Conidia were resuspended in sterile water and the concentration determined using a Neubauer haemocytometer. Viable cell counts were performed by serially diluting the suspension and inoculating onto solid medium to yield between 10 and 100 conidia per plate following 24 h incubation at 37 °C to determine conidia viability. The colony forming units obtained per plate were used to adjust the viable conidial suspensions to the desired inoculum. Fresh conidial suspensions were prepared prior to each independent experiment. All procedures involving manipulation of *A. fumigatus* were performed in a Class II microbiological safety cabinet (Enavir Bio 2+).

##### Generation of *A. fumigatus* Δ*gna1* mutant

PCR primers A & B were designed to amplify a 1189 bp upstream non-coding region of the *GNA1* sequence immediately prior to the ATG start codon and primers C & D, a 1482 bp downstream non-coding region of *GNA1* immediately after the stop codon. Each PCR product was cloned into the pBlueScript $SK^+$ cloning vector (Stratagene). The URA-blaster (8.3 kb) was released by the digestion of plasmid CDA14 with *Hpa*I, separated by agarose gel electrophoresis and extracted using a GeneJET Gel Extraction Kit (Thermo Scientific) according to the manufacturer's instructions. The 8.3 kb DNA fragment was blunt cloned into the site between the up- and downstream non-coding flanking regions of the *GNA1* sequence, to yield the *GNA1* deletion construct pΔ*gna1pyrG+*. The pΔ*gna1pyrG+* construct was transformed into *Escherichia coli* DH5α cells and verified by DNA sequencing (School of Life Sciences, University of Dundee).

To obtain sufficient plasmid DNA, a single colony of *E. coli* DH5α containing pΔ*gna1pyrG+* was inoculated into 5 ml of LB + 50 µg/ml carbenicillin (LB+Amp) medium and incubated at 37 °C under constant agitation (230 rpm) for 8 h. From this starter culture, 200 µl were inoculated into 150 ml of LB+Amp medium (1 in 750 dilution) and incubated as above for 16 h. The cells were harvested by centrifugation for 15 min at 4 °C and the pellet purified using a HiSpeed Plasmid Maxi Kit (Qiagen) according to the manufacturer's instructions. DNA concentration was measured with a NanoDrop Spectrophotometer (Thermo Scientific) and yielded approximately 200 µg of circular plasmid DNA (1 ml at 200 ng/µl). Engineered *Not*I restriction sites facilitated release of an 11 kb linear DNA fragment of pΔ*gna1pyrG+* from the vector backbone. Each *A. fumigatus* protoplast transformation required a 20 µl aliquot containing 5 µg of linear Δ*gna1pyrG+* DNA.

Approximately 1 x $10^6$ cfu/ml of the auxotrophic *A. fumigatus* strain *Ku80ΔpyrG-* was inoculated into 50 ml of complete liquid medium supplemented with 5 mM uridine and 5 mM uracil (CMU) and incubated at 37 °C for 4.5 h with agitation (200 rpm). Following transition from resting to swollen, the conidia were harvested by centrifugation for 10 min at 4 °C and the culture medium discarded. The conidia were then washed twice in 50 ml of sterile water and centrifuged as above. To digest the cell wall, the conidia were incubated for 2 h at 33 °C (120 rpm) in 30 ml of freshly prepared *A. fumigatus* digest solution containing 300 mg of lysing enzymes from *Trichoderma harzianum* (Glucanex®, Sigma). The protoplasts were centrifuged and washed twice in 20 ml of *A. fumigatus* wash solution. Approximately 0.5 ml of *A. fumigatus* store solution was added to the protoplasts to produce a final concentration of 1 x $10^7$ cfu/ml. Aggregates were removed by filtration with a Miracloth (Millipore). Protoplasts were stored at 4 °C and all transformations completed within 12 h.

For transformation, 5 µg of linear Δ*gna1pyrG+* DNA, 200 µl of protoplasts and 50 µl of PEG solution were combined and incubated for 20 min. Following addition of 1 ml of PEG solution, the protoplast-DNA mixture was gently inverted and incubated for a further 20 min at room temperature. Next, this was added to molten 0.35 % agar and 1 ml aliquots overlain on solid transformation medium. Plates were incubated at 35 to 37 °C and inspected for transformants daily after an initial 48 h incubation. All single colonies representative of individual transformants were inoculated on solid medium and incubated at 37 °C for up to 120 h to identify putative Δ*gna1* mutant candidates that were only viable in the presence of exogenous GlcNAc. At least three rounds of phenotypic screening were performed per prospective candidate followed by PCR screening. Approximately 25 ng/ul of genomic DNA was required per PCR reaction using primer pairs to distinguish mutant from wild type (Figure S1).

For Southern Blots, oligonucleotide primers and restriction sites are provided in *Supplemental Experimental Methods*. The PCR DIG Probe Synthesis Kit (Roche) was used to simultaneously amplify and label the probes according to the manufacturer's instructions using 10 pg of plasmid DNA template. The rDIG labelled products were used as probes. Samples of genomic DNA (10 µg) from *A. fumigatus* wild-type (YJ-407) and putative Δ*gna1* mutants were separated on a 0.85 % agarose gel, transferred to a nylon membrane (Roche)[3,4] and the DNA fixed to the membrane by UV crosslinking. The membrane was hybridised overnight in 3.5 ml of DIG Easy Hyb solution (Roche) at 48 °C with the DIG-labelled probes (14 µl of the downstream and 2 µl of the *pyrG* probe following by washed twice in low stringency conditions (25 °C, 5 min, 2× SSC buffer with 0.1 % SDS) and twice in high stringency conditions (68 °C, 15 min, 0.5× SSC with 0.1 % SDS). The bound probe was detected by enzyme immunoassay using the DIG Detection Kit (Roche) as per the manufacturer's instructions.

**Generation of an *A. fumigatus GNA1*+6H::pyrG+ reconstituted strain**

**Design and construction of plasmid *GNA1*+6H**

Procedures followed those for generation of the Δ*gna1* strain, with key reagents in Table S1 and *Supplemental Experimental Methods*. All single colonies representative of individual transformants were inoculated on solid medium and incubated at 37 °C for 24 - 48 h to identify putative auxotrophic *A. fumigatus GNA1*+6H::*pyrG*- reconstitutes that were only viable in the presence of exogenous uridine and uracil. At least three rounds of phenotypic screening were performed per prospective candidate to assess stability of the growth phenotype. Any candidates that grew on CM were discarded from subsequent analysis due to ectopic retention of the *pyrG*+ selection marker. PCR screening and Southern Blot verification was performed as for the Δ*gna1* mutant, with primers listed in *Supplemental Experimental Methods*.

To permit *in vivo* applications by restoring uridine and uracil autotrophy, the reconstituted strain (*GNA1*+6H::*pyrG*-) was complemented with *pyrG* by random integration. Primers were used to amplify a 1.9 kb portion of the *A. fumigatus pyrG* gene from pXDRFP4 (Fungal Genetics Stock Center, Kansas City, Missouri USA)[5]. *A. fumigatus GNA1*+6H*::pyrG*- protoplasts were transformed with a 20 µl aliquot containing a 5 µg fragment of *AfpyrG* (see relevant information in *Supplemental Experimental Methods*). The transformant colonies were not amenable to phenotypic screening as viability on CM was indicative of restoration of the wild-type growth phenotype. Eight individual transformants were randomly selected for PCR screening to verify integration of *pyrG* and the *GNA1* promoter and coding region remained intact and not subject to ectopic integration.

***In vitro* phenotypic analysis of *A. fumigatus* Δ*gna1* strains**

***Analysis of cell wall architecture* by TEM and SEM**

For analysis of the cell wall architecture, conidia and mycelia were grown on solid medium, fixed, processed and examined using electron microscopy. For SEM, conidia were fixed in 2.5 % glutaraldehyde for 24 h, washed twice for 15 mins in 0.2 M piperazine-N,N'-bis(2-ethanesulfonic acid) (PIPES) pH 7.2 buffer and immersed in PBS for 16 - 18 h. The specimens were dehydrated through a graded ethanol series and transferred to 100 % acetone for 15 mins. A critical point dryer (CPD 030, Bal-Tec) was used to dry and preserve the delicate surface structures. The specimens were mounted on aluminium stubs using carbon adhesive tabs, coated with 40 nm gold/palladium using a High-Resolution Sputter Coater (208 HR, Cressington Scientific) and examined under a scanning electron microscope (Hitachi S-4700) operating at an accelerating voltage of 5 kV. For TEM, hyphae were fixed in 2.5 % glutaraldehyde for up to 24 h, washed twice for 15 mins in 0.2 M PIPES pH 7.2 buffer and immersed in 1 % aqueous osmium tetroxide for 1 h. The specimens were washed twice for 15 min in distilled water before fixation/staining in 3 % aqueous uranyl acetate for 16 - 18 h. Following washing in distilled water (as above), the specimens were dehydrated through a graded ethanol series and immersed in propylene oxide twice for 15 min each. A 50 % mixture of Durcupan resin and propylene oxide was infiltrated into the specimens for 24 h on a rotary mixer at ambient temperature. After infiltration for a further 24 h in 100 % Durcupan resin, the specimens were polymerised at 60 °C for 24 h. Sections were cut using a microtome (Ultracut UCT, Leica), mounted on pioloform coated copper grids, post stained with 3 % aqueous uranyl acetate and Reynold's lead citrate and examined using a transmission electron microscope (JEOL-1200 EX) operating at an accelerating voltage of 80 kV.

**PCR primers to generate and verify the *A. fumigatus* Δ*gna1* mutant and reconstituted strains.** The reverse complement sequence is provided for all antisense (R) primers. **A to D**, primers for amplification and cloning of *GNA1* upstream and downstream non-coding flanking regions for homologous recombination. **P1 to P10**, primers for verification of a Δ*gna1* mutant. Up-up and Down-down denote regions extending beyond the upstream and downstream regions of homologous recombination. **Pi to Piv**, primers for the generation of Southern Blot probes. **1 to 6**, primers for amplification and ligation to generate p*GNA1*+6H*::pyr*G- (auxotrophic reconstituted strain). **P11 to P18**, primers to differentiate the parental, Δ*gna1* mutant and reconstituted *pyrG*- strains. **P19 to P20**, primers for amplification of *A. fumigatus pyrG* open reading frame.

| Primer ID | Description | Sequence (5′ to 3′) | Product size (bp) |
|---|---|---|---|
| A | *GNA1* upstream flanking (F) | AAACTCGAGGCGGCCGCCTGCTTCTTCTTTCCATTCATCCCTCC | 1189 |
| B | *GNA1* upstream flanking (R) | CTCACTCTCGAACTCAACGATATCTTTGGGATTGATCCTGTCGGTAACGG | |
| C | *GNA1* downstream flanking (F) | CAGGATCAATCCCAAAGATATCGTTGAGTTCGAGAGTGAGCTGGACAG | 1484 |
| D | Downstream flanking (R) | TTTTCTAGAGCGGCCGCTGAAGTTTTTGATCTCCACGTTCCTGC | |
| P1 | *GNA1* native (F) | ATGACCAACGCAACCATTGCTCCGA | 644 |
| P2 | *GNA1* native (R) | TCAGTAGTAGTGCGCCATCTCCAAC | |
| P3 | *GNA1* upstream 366 (F) | GAGTAACTGTTCGGTGATGC | 1510 |
| P4 | *GNA1* downstream 500 (R) | TGATTATGCTTGCTGTATTGC | |
| P5 | Up-Up 171 (F) | GCTTGCTGAACCTGGGTACT | 1466 |
| P6 | *neo*-up 195 (R) | AGTAATCGCAACATCCGCAT | |
| P7 | *An pyrG* 3794 (F) | GCCATTTTCTCTACGCAAGAGT | 1355 |
| P8 | *An pyrG* 5149 (R) | GCGATGATTTTACTGAGCTTGC | |
| P9 | *neo*-down 588 (F) | CCATCACGAGATTTCGATTC | 2203 |
| P10 | Down-down 115 (R) | CAACTTACCATCTCGATCCG | |
| Pi | Probe 1 (F) | GCCATTTTCTCTACGCAAGAGT | 1246 |
| Pii | Probe 1(R) | CGTCTCTGGATTTACGAATCAG | |
| Piii | Probe 2 (F) | CAGGATCAATCCCAAAGATATCGTTGAGTTCGAGAGTGAGCTGGACAG | 1484 |
| Piv | Probe 2 (R) | TTTTCTAGAGCGGCCGCTGAAGTTTTTGATCTCCACGTTCCTGC | |
| 1 | Upstream (F) | AAACTCGAGCTGCTTCTTCTTTCCATTCATCCCTCC | 1189 |
| 2 | Upstream (R) | GTCGGAGCAATGGTTGCGTTGGTCATTTTGGGATTGATCCTGTCGGTAACGG | |
| 3 | *GNA1* (F) | CCGTTACCGACAGGATCAATCCCAAAATGACCAACGCAACCATTGCTCCGAC | 644 |
| 4 | Upstream_*GNA1* (R) | GAATCTTTTACCAGATCGGAAGCAATTGGTCAGTAGTAGTGCGCCATCTCCAACC | |
| 5 | Downstream (F) | CTGGGTTGGAGATGGCGCACTACTACCATCACCATCATCACCACTGATCTAGACCCGGGGTTGAGTTCGAGAGTGAGCTGG | 1551 |
| 6 | Downstream (R) | GAATCTTTTACCAGATCGGAAGCAATTGGTTTGCGGCCGCTGAAGTTTTT GATC | |
| P11 | *GNA1* promoter (F) | CGTTACCGACAGGATCAATC | 676 |
| P12 | *GNA1* stop codon flank (R) | ACTCAACTCAGTAGTAGTGC | |

| | | | |
|---|---|---|---|
| P13 | *GNA1* promoter (F) | CGTTACCGACAGGATCAATC | 681 |
| P14 | *GNA1* linker (R) | GTGATGATGGTGATGGTAGT | |
| P15 | Up-Up 171 (F) | GCTTGCTGAACCTGGGTACT | 1936 |
| P16 | *GNA1* linker (R) | GTGATGATGGTGATGGTAGT | |
| P17 | *GNA1* linker (F) | ACTACCATCACCATCATCAC | 1664 |
| P18 | Down-down 115 (R) | CAACTTACCATCTCGATCCG | |
| P19 | *Af pyrG* (F) | GCTCGAGCATGCATCTAGAG | 1913 |
| P20 | *Af pyrG* (R) | CTGTCTGAGAGGAGGCACTG | |

**Solutions for *A. fumigatus* protoplast transformation.**

| Solution | Ingredients and method (per 100 ml) |
|---|---|
| Cell wall digest | $(NH_4)_2SO_4$ (Ammonium sulphate) ........ 5.280 g<br>Potassium citrate........ 1.620 g<br>Yeast extract........ 0.500 g<br>Sucrose........ 0.500 g<br>$MgSO_4$ ........ 0.246 g<br>Using concentrated HCl adjust to pH 6.0. |
| Protoplast wash | $(NH_4)_2SO_4$ (Ammonium sulphate) ........ 5.28 g<br>Sucrose........ 1.00 g<br>Potassium citrate........ 1.62 g<br>Using concentrated HCl adjust to pH 6.0. |
| Protoplast store | KCl ........ 4.440 g<br>$CaCl_2$........ 0.555 g<br>MOPS ........ 0.200 g<br>Using NaOH adjust to pH 6.0. |
| PEG for transformation | PEG 6000........ 25.000 g<br>KCl ........ 4.440 g<br>$CaCl_2$........ 0.555 g<br>Tris........ 0.121 g<br>Using concentrated HCl adjust to pH 7.5. |

**Medium for protoplast transformation and phenotypic screening.** MM, minimal medium; NGMM, non-glucose minimal medium; CM, complete medium; NGCM, non-glucose complete medium; 90 mM GlcNAc; N/A, not applicable.

| | ***A. fumigatus* strain** | | |
|---|---|---|---|
| | Δ*gna1*<br>(mutant) | *GNA1*+6H*pyrG*-<br>(Reconstituted - auxotrophic) | *GNA1*+6H*pyrG*+<br>(Reconstituted) |
| **Transformation** | NGMM + 1 M sorbitol + 90 mM GlcNAc | MM + 1 M sorbitol + 5 mM uridine + 5 mM uracil | MM + 1 M sorbitol |
| **Positive phenotypic growth control** | NGCM +<br>90 mM GlcNAc | CM +<br>5 mM uridine + 5 mM uracil | CM |
| **Negative phenotypic growth control** | CM | CM | N/A |

##### *Af*Gna1 fragment screen and Biolayer Interferometry (BLI) binding affinity measurements

To identify a potential fragment binder of *Af*Gna1, the fragment library of the University of Dundee Drug Discovery Unit (DDU) comprising of 652 structurally diverse compounds was screened using biolayer interferometry (BLI). To do this *Af*Gna1 was first buffer exchange into biotinylation buffer (25 mM Hepes, 150 mM NaCl, 0.5 mM TCEP pH 7.5) and biotinylated by incubation with biotin reagent (EZ-Link® NHS-Peg$_4$-Biotin,Thermo Scientific) in a 1:1 molar ratio at 4 °C for 2 hrs. Excess biotin reagent was removed by buffer exchange using a 2 ml Zeba spin desalting column (Thermo Fisher) according to the manufacturer's protocol.

BLI experiments were performed using the Octet Red 384 system (Forte Bio, Menlo Park, CA USA). Briefly super streptavidin (SSA) biosensors were hydrated in assay buffer (25 mM Hepes, 150 mM NaCl, 0.5 mM TCEP pH 7.5. 1 mM CoA) for a minimum of 5 min followed by protein immobilisation to sensors test to determine the best optimum concentration for the experiment. The optimum concentration for this experiment was ~25 µg/ml. For the actual fragment screening, *Af*Gna1 was immobilised on 16 SSA biosensors in parallel for 900s, and free streptavidin sites on the sensors were blocked with biocytin for 120 s, another set of 16 SSA biosensors were blocked with biocytin also for 120 s and later washed in the assay buffer (25 mM Hepes, 150 mM NaCl, 0.5 mM TCEP pH 7.5. 1 mM CoA) this serve as the control for reference sensors. The fragment screening consisted of a single point assay of three 60 s steps: (a) baseline in assay buffer, (b) association in each fragment at 200 µM and (c) dissociation in assay buffer. The entire screen was repeated with the control (reference) sensors. Data were processed and visualised using Octet software and the response rate for each fragment was determined by subtracting the baseline and reference responses from those obtained in the presence of Gna1. Primary hits were defined as fragments with a response rate greater than three robust standard deviation units above the median.

Confirmation of each primary hit was performed with a follow-up experiment using a 6-point concentration dependent series commencing at 500 µM in 3-fold serial dilutions, unless described otherwise. The assay was performed as described earlier with the dissociation step increased to 120 s using biosensors immobilised with Gna1 followed by blocked streptavidin control biosensors. Binding isotherms were fit using Octet software to determine the dissociation constant ($K_d$). The $K_d$ value was double referenced by applying global, steady state and partial fits (where appropriate). Fragments of interest were identified by visual inspection of the sensorgrams and the steady state fit.

##### Isothermal Titration Calorimetry

ITC was conducted on a MicroCal PEAQ-ITC instrument (Malvern). *Af*GNA1 was dialysed against argon-purged ITC buffer (20 mM HEPES, 100 mM NaCl, 0.25 mM TCEP, pH 7.5). The sample cell contained *Af*Gna1 (50 µM); The syringe contained fragment hit **1** (1000 µM); In all, 19 injections were delivered (0.8 s addition time, interval 300 s). The stirrer speed was set to 750 rpm, and binding experiments were carried out at 25 °C. Water was used in the reference cell, and titrations into buffer were carried out to assess the enthalpies of dilution. The data was smoothed with the simple arithmetic function in the MicroCal PEAQ-ITC Analysis Software (Malvern).

##### General synthetic and analytical procedures

Chemicals were purchased from Sigma-Aldrich, TCI UK, Fluorochem, Alfa Aesar, Carbosynth and WuXi Apptec and used without further purification. All laboratory-reagent grade, analytical grade and anhydrous solvents were purchased from Sigma-Aldrich, Fisher Scientific and Acros Organics. Air- and moisture-sensitive reactions were carried out under an inert atmosphere of argon in dried glassware. Thin-layer chromatography (TLC) was performed on precoated TLC plates (POLYGRAM® SIL G/UV$_{254}$, Macherey-Nagel). Developed plates were dried and analysed under a UV lamp (UV254/365 nm). Ultra-pure water and ddH$_2$O was obtained from a Millipore milliQ (MQ) Advantage system. Small molecule LCMS was carried out using electrospray ionization (ESI) on an Agilent Technologies 1200 single quadrupole LC-MS system fitted with a Max-Light Cartridge flow cell coupled to a 6130 Quadrupole spectrometer. An Agilent ZORBAX Eclipse Plus C18, 4.6 x 100 mm, 3.5 µm column was used for separations. Variable wavelengths were used and MS acquisitions were carried out in positive and negative ion modes. Intact protein mass was carried out using the same Agilent instrument using an Agilent ZORBAX 300SB-C3 5um, 2.1 x 150mm column, unless otherwise stated. Protein MS acquisition was carried out in positive ion mode and total protein masses were calculated by deconvolution within the MS Chemstation software (Agilent Technologies). The LCMS solvent system consisted of 0.05 % trifluoroacetic acid (TFA) in $H_2O$ as buffer A, and 0.04 % TFA acid in acetonitrile (ACN) as buffer B. Protein UV absorbance was monitored at 214 nm, 254 nm and 280 nm.

NMR spectra were recorded on a Bruker Avance II 500 MHz spectrometer or a Bruker Ascend 400 MHz spectrometer at room temperature. Chemical shifts (δ) are expressed in ppm recorded using the residual solvent as the internal reference in all cases. The following abbreviations are used to indicate the signal multiplicity: s (singlet), bs (broad singlet), d (doublet), t (triplet), q (quartet), dd (doublet of doublets), m (multiplet). High resolution mass spectrometry (HRMS) was carried out using electrospray ionization (ESI) on a Bruker Daltonics MicroTOF mass spectrometer.

**Supplemental Fragment derivatives**

The synthesis of 1,2,4-triazole fragments highlighted in FigureS2d was achieved using reported literature methods (unless commercially available). In brief, generation of the 1,2,4-triazole scaffold is achieved in two steps with a suitable amide-containing precursor by reaction with *N,N*-dimethylformamide dimethylacetal (DMFDMA) followed by subsequent ring closure with hydrazine[6]. Where possible an amide-containing precursor was used to expedite synthesis, however, if not commercially practical, a carboxylic acid[7,8] or nitrile precursor[9] was used as an amide synthon for use in subsequent 1,2,4-triazole-formation reactions. The anilino functionality present on the aryl ring (**S11-S15**, **S20**), was introduced by nitro-group reduction of suitably functionalised starting materials[10,11]. The pyrrole ring (**S22**) was introduced using a suitable vinylbenzene precursor according to literature procedure[12].

**S8**

$^{1}$H NMR (500 MHz, DMSO-D6): $\delta$ = 14.23 (bs, 1H), 8.47 (bs, 1H), 8.01 (dd, J = 1.8 Hz, 7.7 Hz, 1 H), 7.54 - 7.46 (m, 1H), 7.30 -7.39 (m, 2H).
$^{19}$F NMR (470 MHz, DMSO-D6): $\delta$ = -112.94 (s).
$^{13}$C NMR (101 MHz, DMSO-D6): $\delta$ = 160.76, 158.76, 143.89 - 149.75 (bs), 131.68, 130.35, 125.16, 116.99, 116.82.
LCMS: Theo. m/z 163.1 Da; obs. m/z 164.1Da [M+H].

**S9**

$^{1}$H NMR (500 MHz, DMSO-D6): $\delta$ = 14.25 (bs, 1H), 8.61 (bs, 1H), 7.88 (d, J = 8.1 Hz, 1H), 7.74 - 7.84 (m, 2H), 7.66 - 7.72 (m, 1H).
$^{19}$F NMR (470 MHz, DMSO-D6): $\delta$ = -57.30 (s).
$^{13}$C NMR (101 MHz, DMSO-D6-D4): $\delta$ = 143.96 (bs), 132.77, 132.31, 129.66 - 130.41 (bs), 126.96 (m), 125.38, 123.20.
LCMS: Theo. m/z 213.1 Da; obs. m/z 214.1Da [M+H], 194.1 Da [M-F].

**S10**

$^{1}$H NMR (500 MHz, $CDCl_3$): $\delta$ = 8.16 (s, 1H), 7.68 (dd, J = 1.4 Hz, 7.7 Hz, 1H), 7.20 - 7.40 (m, 4H), 2.54 (s, 3H).
$^{13}$C NMR (101 MHz, $CDCl_3$): $\delta$ = 148.12, 137.17, 131.41, 129.95, 129.43, 126.15, 21.07.
LCMS: Theo. m/z 159.1 Da; obs. m/z 160.1 Da [M+H].

**S11**

$^{1}$H NMR (500 MHz, MeOH-D4): $\delta$ = 7.95 (bs, 1H), 7.79 (d, J = 8.5 Hz, 1H), 6.45 (d, J = 1.3 Hz, 1H), 6.40 (dd, J = 1.6 Hz, 8.3 Hz, 1H), 3.95 (s, 3H).
$^{13}$C NMR (101 MHz, MeOH-D4): $\delta$ = 159.98, 153.83, 149.59 - 151.07 (bs), 131.31, 108.44, 98.07, 55.85.
LCMS: Theo. m/z 190.1 Da; obs. m/z 191.1 Da [M+H].

**S12**

$^{1}$H NMR (500 MHz, MeOH-D4): $\delta$ = 7.85 (bs, 1H), 7.52 (dd, J = 8.4 Hz, 8.5 Hz, 1H), 6.41 (dd, J = 2.0 Hz, 8.6 Hz, 1 H), 6.33 (dd, J = 2.0 Hz, 13.7 Hz, 1 H).
$^{19}$F NMR (470 MHz, MeOH-D4): $\delta$ = -76.93 (s).
$^{13}$C NMR (101 MHz, MeOH-D4): $\delta$ = 162.36, 160.40, 149.52 - 150.79 (bs), 129.88, 110.42, 100.11.
LCMS: Theo. m/z 178.1 Da; obs. m/z 179.1 Da [M+H].

**S13**

$^{1}$H NMR (500 MHz, MeOH-D4): $\delta$ = 13.91 (bs, 1H), 7.38 (bs, 1H), 7.0 (s, 1H), 6.83 (d, J = 8.0 Hz, 1H), 5.81 (bs, 2H).
$^{13}$C NMR (101 MHz, MeOH-D4): $\delta$ = 133.21, 116.47, 111.28, resonances absent.
$^{19}$F NMR (470 MHz, MeOH-D4): $\delta$ = -57.72 (s).
LCMS: Theo. m/z 228.1 Da; obs. m/z 229.1 Da [M+H].

**S14**

$^{1}$H NMR (500 MHz, MeOH-D4): $\delta$ = 7.30 (bs, 1H), 6.64 (d, J = 1.5 Hz, 1H), 6.60 (dd, J = 2.1 Hz, 8.2 Hz, 1H), 2.36 (s, 3H).
$^{13}$C NMR (101 MHz, MeOH-D4): $\delta$ = 149.46 - 151.20 (bs), 137.91, 130.20, 116.52, 112.08, 39.07, 19.26.
LCMS: Theo. m/z 174.1 Da; obs. m/z 175.1 Da [M+H].

**S15**

$^{1}$H NMR (500 MHz, MeOH-D4): $\delta$ = 7.92 - 8.38 (bs, 1H), 7.43 (d, J = 8.4 Hz, 1H), 6.80 (d, J = 2.2 Hz, 1H), 6.67 (dd, J = 2.3 Hz, 8.5 Hz, 1H).
$^{13}$C NMR (101 MHz, MeOH-D4): $\delta$ = 149.08 - 151.33 (bs), 132.96, 131.72, 131.33, 114.49, 112.67, 111.27.

LCMS: Theo. m/z 194.1 Da; obs. m/z 195.0, 197.0 Da [M+H].

**S19**

$^{1}$H NMR (500 MHz, MeOH-D4): $\delta$ = 8.45 - 8.64 (bs, 1H), 7.49 - 7.58 (m, 3H).
$^{13}$C NMR (101 MHz, MeOH-D4): $\delta$ = 144.72 - 149.65 (bs), 137.28, 133.17, 129.36, resonances absent.
LCMS: Theo. m/z 213.0 Da; obs. m/z 214.0, 216.0 Da [M+H].

**S20**

$^{1}$H NMR (500 MHz, MeOH-D4): $\delta$ = 8.47 (s, 1H), 6.76 (s, 2H).
$^{13}$C NMR (101 MHz, MeOH-D4): $\delta$ = 154.46, 153.27, 147.91, 137.17, 114.59, 114.09.
LCMS: Theo. m/z 228.0 Da; obs. m/z 229.0, 231.0 Da [M+H].

**S21**

$^{1}$H NMR (500 MHz, DMSO-D6): $\delta$ = 8.39 (bs, 1H), 7.29 (dd, J = 7.5 Hz, 7.6 Hz, 1H), 7.16 (d, J = 7.7 Hz, 1H), 2.05 (s, 6H).
$^{13}$C NMR (101 MHz, DMSO-D6): $\delta$ = 146.77, 142.48, 138.44, 136.73, 136.40, 29.15.
LCMS: Theo. m/z 173.1 Da; obs. m/z 174.1 Da [M+H].

**S22**

$^{1}$H NMR (500 MHz, MeOH-D4): $\delta$ = 7.36 (d, J = 8.0 Hz, 2H), 7.13 (dd, J = 7.8 Hz, 7.9 Hz, 1H), 6.77 - 6.81 (m, 2H), 6.16 (dd, J = 1.6 Hz, 1.6 Hz, 1H).
$^{13}$C NMR (101 MHz, MeOH-D4): $\delta$ = 136.90, 136.73, 129.16, 128.85, 119.36, 118.68, 118.04, 110.55.
LCMS: Theo. m/z 211.0 Da; obs. m/z 212.0 Da [M+H].

**Commercially available fragments.** Refer to Figure S2.

| *Cpd.* | *Name* | *Vendor* | *Vendor code* |
|---|---|---|---|
| **S4** | 3-phenyl-1,2,4-triazole | Sigma | S294217 |
| **S5** | 4-(1,2,4-triazol-5-yl)benzoic acid | Enamine | EN300-139972 |
| **S6** | 4-(1,2,4-triazol-3-yl)phenol | Enamine | EN300-77930 |
| **S7** | 4-(1,2,4-triazol-3-yl)aniline | Enamine | EN300-74655 |
| **S16** | 3-(2-chlorophenyl)-4-methyl-1,2,4-triazole | Enamine | EN300-142445 |
| **S17** | 3,5-diphenyl-1,2,4-triazole | Enamine | EN300-107143 |
| **S18** | Ethyl 5-phenyl-1,2,4-triazole-3-carboxylate | Key Organics | 4M-441S |
| **S23** | 3-(3-chlorophenyl)-1,2,4-triazole | Princeton Bio | PBMR111005 |
| **S24** | 3-(1,2,4-triazole-3-yl)aniline | Enamine | EN300-76519 |

**NMR data**

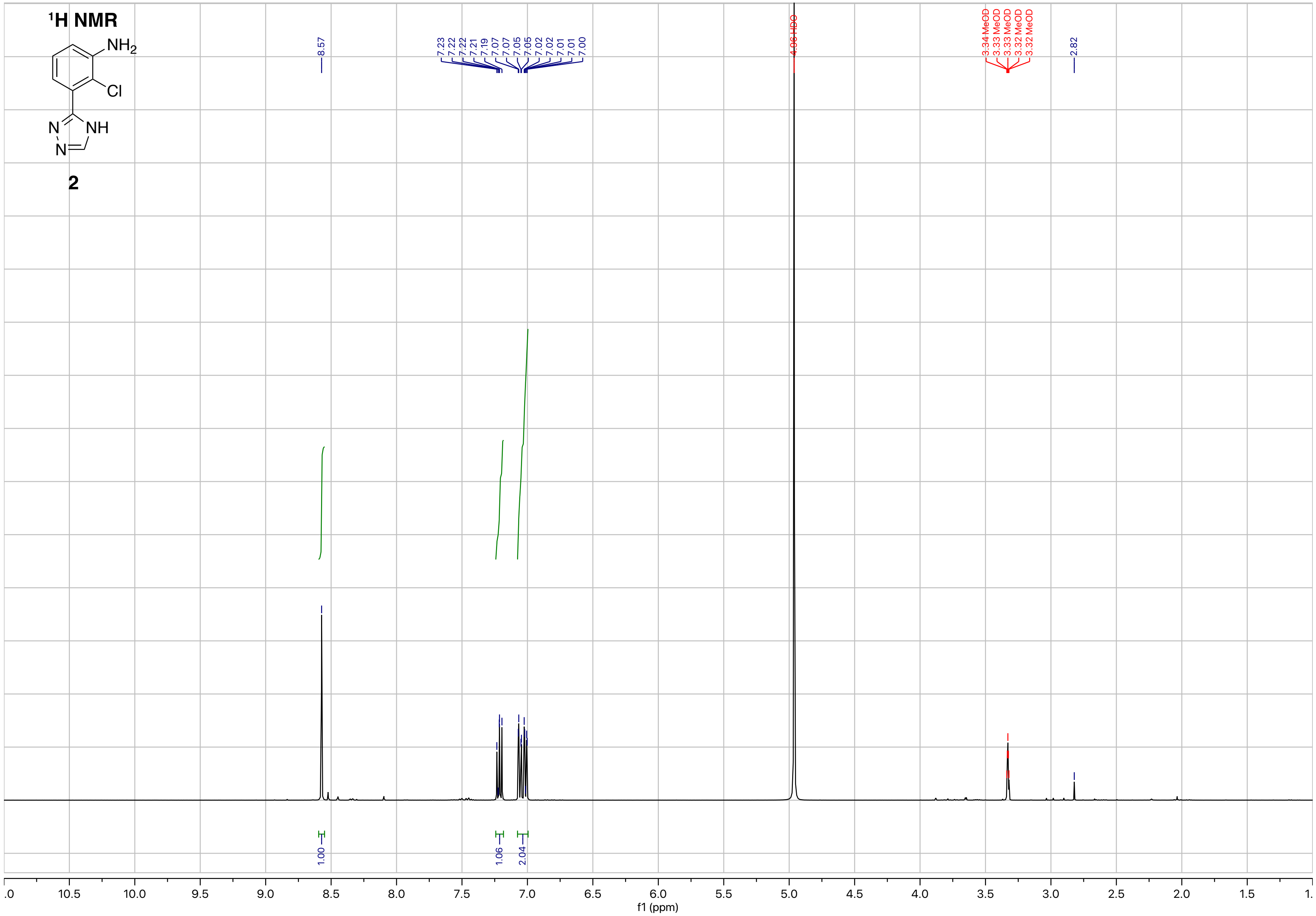

¹H NMR
NH2
Cl
N
NH
N
2
8.57
7.23
7.22
7.22
7.21
7.19
7.07
7.07
7.05
7.05
7.02
7.02
7.01
7.01
7.00
4.96 HDO
3.34 MeOD
3.33 MeOD
3.33 MeOD
3.32 MeOD
3.32 MeOD
2.82
1.00
1.06
2.04
f1 (ppm)

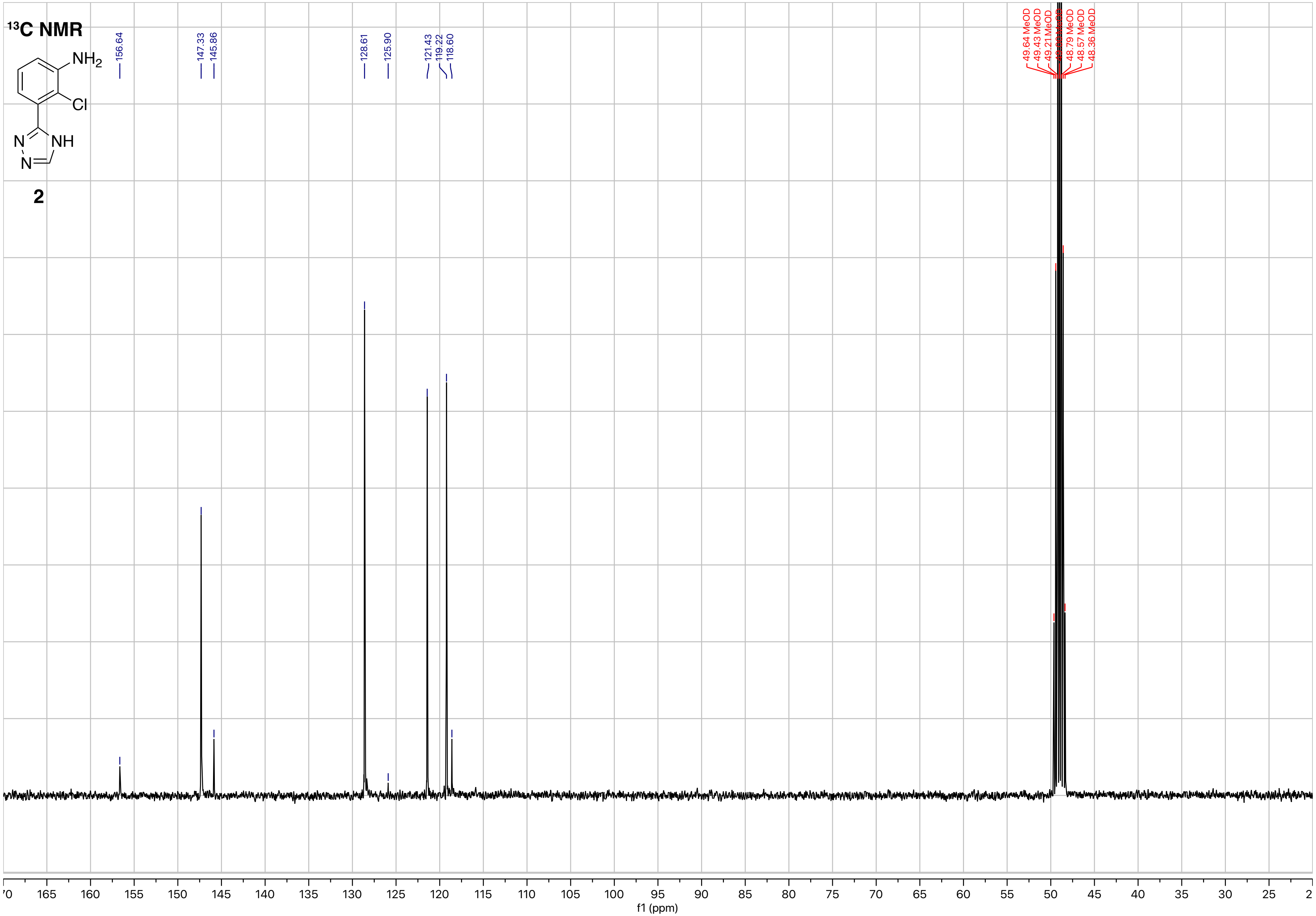

¹³C NMR
NH2
Cl
N
NH
N
2
156.64
147.33
145.86
128.61
125.90
121.43
119.22
118.60
49.64 MeOD
49.43 MeOD
49.21 MeOD
48.79 MeOD
48.57 MeOD
48.36 MeOD
165
160
155
150
145
140
135
130
125
120
115
110
105
100
95
90
85
80
75
70
65
60
55
50
45
40
35
30
25
f1 (ppm)

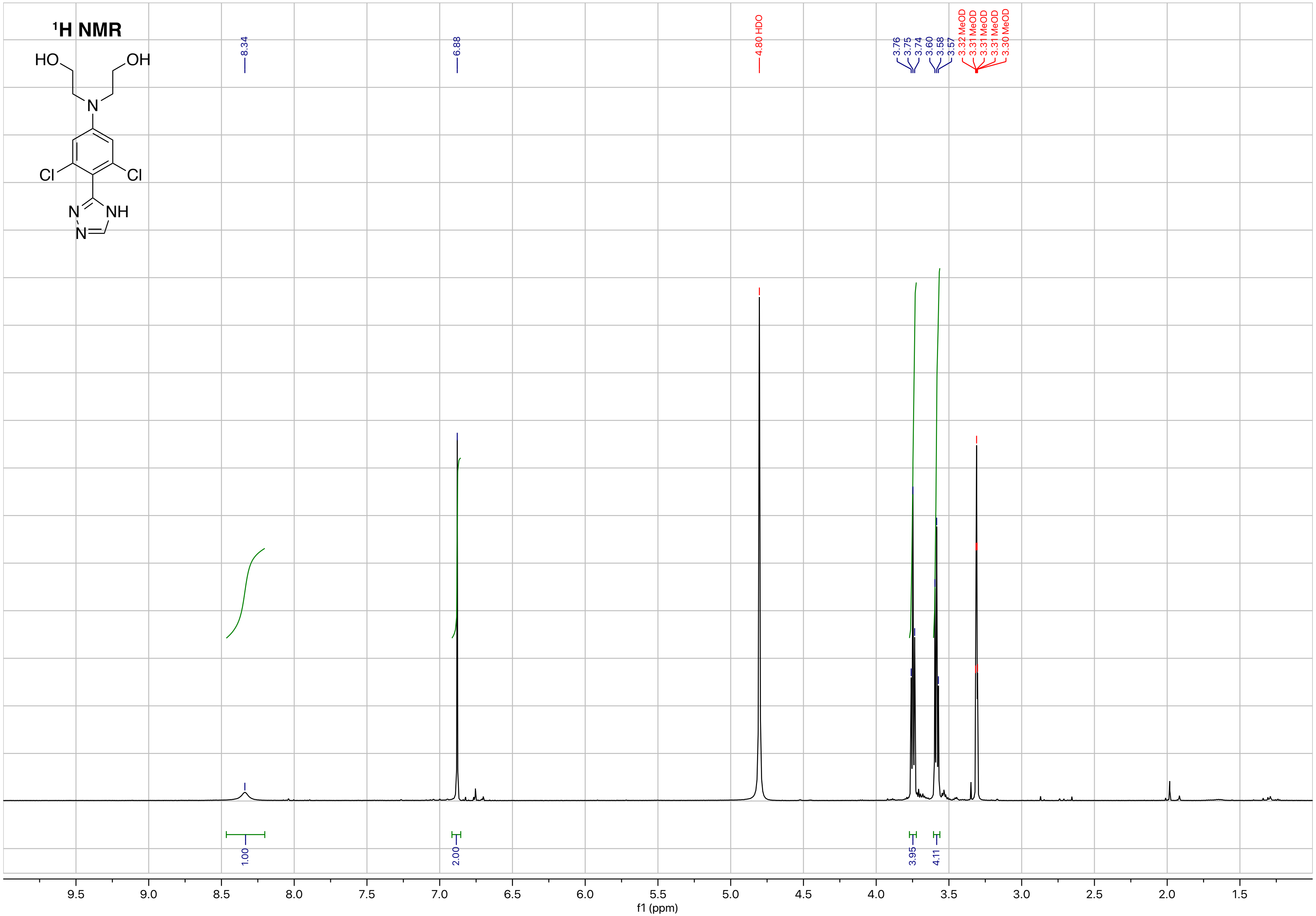

1H NMR
HO
OH
N
Cl
Cl
N
NH
N
8.34
6.88
4.80 HDO
3.76
3.75
3.74
3.60
3.58
3.57
3.32 MeOD
3.31 MeOD
3.31 MeOD
3.31 MeOD
3.30 MeOD
1.00
2.00
3.95
4.11
9.5
9.0
8.5
8.0
7.5
7.0
6.5
6.0
5.5
5.0
4.5
4.0
3.5
3.0
2.5
2.0
1.5
f1 (ppm)

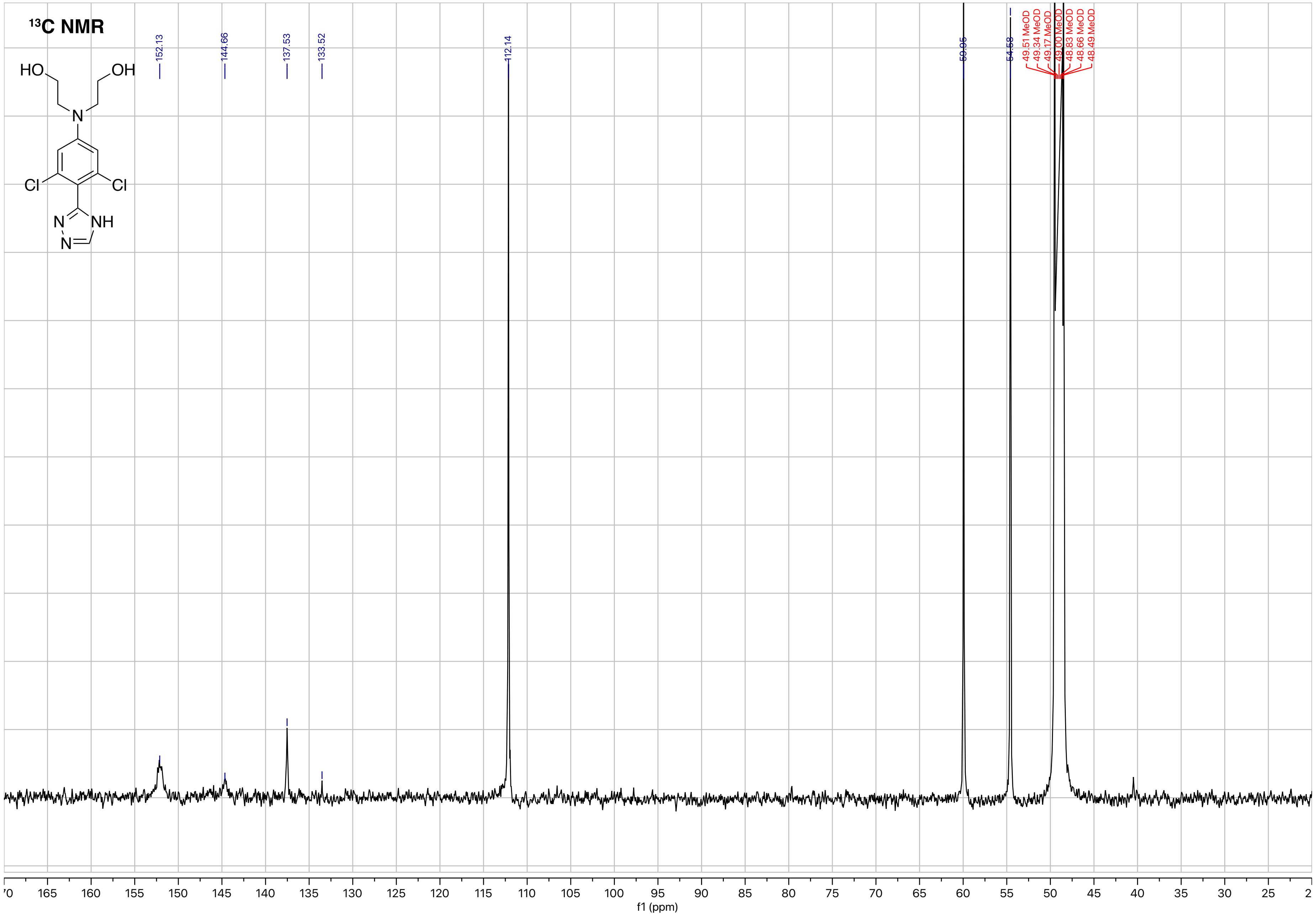

13C NMR
152.13
144.66
137.53
133.52
112.14
59.95
54.58
49.51 MeOD
49.34 MeOD
49.17 MeOD
49.00 MeOD
48.83 MeOD
48.66 MeOD
48.49 MeOD
HO
OH
N
Cl
Cl
N
NH
N
f1 (ppm)

**Supplemental References**


1. Robert, X. & Gouet, P. Deciphering key features in protein structures with the new ENDscript server. *Nucleic Acids Res.* **42**, W320–W324 (2014).
2. Chung, C. C. *et al.* A fluorescence-based thiol quantification assay for ultra-high-throughput screening for inhibitors of coenzyme A production. *Assay Drug Dev. Technol.* **6**, 361–374 (2008).
3. Brown, T. Analysis of RNA by Northern and Slot-Blot Hybridization. in *Current Protocols in Immunology* (2001). doi:10.1002/0471142735.im1012s07
4. Sambrook, J., Fritsch, E. F. & Maniatis, T. *Molecular Cloning a Laboratory Manual Second Edition Vols. 1 2 and 3. Sambrook J E F Fritsch and T Maniatis Molecular Cloning A Laboratory Manual Second Edition Vols 1 2 and 3 XxxixPagination VariesVol 1 XxxiiiPagination VariesVol 2 XxxiiPagination VariesVol 3 Cold Spring Harbor Laboratory Press* (1989).
5. McCluskey, K., Wiest, A. & Plamann, M. The fungal genetics stock center: A repository for 50 years of fungal genetics research. *Journal of Biosciences* **35**, 119–126 (2010).
6. May, C., Sun, Y., Wolmershäuser, G. & Thiel, W. R. Synthetic access to a novel binaphthyl ligand bearing a phosphine and a triazole donor site. *Zeitschrift fur Naturforsch. - Sect. B J. Chem. Sci.* **64**, 1438–1448 (2009).
7. Michel, A. D. & Walter, D. S. Preparations of prolinamide p2x7 modulators and their combinations with other therapeutic agents. *PCT Int. Appl.* 136pp. (2009).
8. Abe, T., Takahashi, Y., Matsubara, Y. & Yamada, K. An Ullmann N-arylation/2-amidation cascade by self-relay copper catalysis: one-pot synthesis of indolo[1,2-a]quinazolinones. *Org. Chem. Front.* **4**, 2124–2127 (2017).
9. Katritzky, A. R., Pilarski, B. & Urogdi, L. Efficient conversion of nitriles to amides with basic hydrogen peroxide in dimethyl sulfoxide. *Synthesis (Stuttg).* 949–951 (1989). doi:10.1055/s-1989-27441
10. Bellamy, F. D. & Ou, K. Selective reduction of aromatic nitro compounds with stannous chloride in non acidic and non aqueous medium. *Tetrahedron Lett.* **25**, 839–842 (1984).
11. Kaplánek, R. & Krchňák, V. Fast and effective reduction of nitroarenes by sodium dithionite under PTC conditions: Application in solid-phase synthesis. *Tetrahedron Lett.* **54**, 2600–2603 (2013).
12. Smith, N. D., Huang, D. & Cosford, N. D. P. One-step synthesis of 3-aryl- and 3,4-diaryl-(1H)-pyrroles using tosylmethyl isocyanide (TOSMIC). *Org. Lett.* **4**, 3537–3539 (2002).
